# Polarization-engineered aberration-resilient light sheet microscopy

**DOI:** 10.64898/2026.05.11.724351

**Authors:** Yuqing Qiu, Juncheng Zhang, Christopher R. Warren, Sara Kacmoli, Vanessa Gonzalez, Cullen B. Young, Mengyang J. Li, Fei Liu, Kevin Keomanee-Dizon, Rebecca D. Burdine, Tian-Ming Fu

## Abstract

Light sheet fluorescence microscopy enables volumetric imaging with high imaging speed, optical sectioning capability, and reduced photobleaching and phototoxicity, and has become a workhorse in bioimaging. However, widely adopted Gaussian light sheets face an inherent trade-off between axial resolution and field-of-view due to diffraction. State-of-the-art ‘nondiffracting’ light sheets—including Bessel beam, Airy beam, and lattice light sheet—alleviate this trade-off but suffer from optical aberrations that compromise performance with increasing imaging depth. While the integration of adaptive optics offers a promising solution, such integrated systems are typically complex, expensive, and slow due to the need for serial mapping and correction of spatially varying aberrations across the specimen. Here, we present polarization-engineered aberration-resilient light sheet (PEARLS), a new class of monochromatic nondiffracting light sheet with temporally invariant profile and robustness to optical aberrations. In comparison with existing light sheets, PEARLS showed significantly reduced photobleaching and enhanced aberration-resilience, permitting imaging of three-dimensional subcellular dynamics in optically complex environments. We applied PEARLS for noninvasive observations of biological dynamics in various living systems, revealing phenotypic diversity across spatial and temporal scales—from rapid membrane dynamics and organelle interactions in cultured cells to coordinated mitosis and cell migrations in developing embryos.

## Main Text

The capability to noninvasively visualize biological dynamics across molecular to multicellular scales deep inside the organisms could offer crucial insights into fundamental biology and broad-reaching implications for human health. In recent years, light sheet fluorescence microscopy (LSFM) has emerged as a workhorse for long-term imaging of living biological specimen with greatly reduced invasiveness^1–14^. LSFM creates a thin sheet of light that selectively illuminates a plane of the specimen and observes the emitted fluorescence perpendicularly (Figure S1A). The light sheet, which is key to the performance of LSFM, offers two unique advantages: First, it provides additional spatial resolution, achieving axial resolution beyond the diffraction limit of the detection objective^3^. Second, compared to widefield and confocal fluorescence microscopy, LSFM minimizes imaging background, photobleaching, and phototoxicity by restricting the illumination to the detection focal plane.

An ideal light sheet profile should simultaneously satisfy three criteria: (i) a propagation-invariant profile that maintains high axial resolution over a large field-of-view (FOV); (ii) a temporally invariant profile with a full duty cycle to minimize peak illumination intensity and phototoxicity; (iii) robustness to optical aberrations when imaging within thick heterogeneous biological tissues. However, existing light sheets inevitably compromise at least one of these criteria. The widely adopted Gaussian light sheets face a fundamental tradeoff between spatial resolution and FOV^14–16:^ its FOV shrinks quadratically as the light sheet thickness decreases (Figure S1B-E). To overcome this fundamental tradeoff, numerous beam-shaping strategies based on so-called “nondiffracting” beams have been explored. However, only two classes of nondiffracting monochromatic beams with one transverse dimension (i.e. light sheet) exist^17^: the cosine standing wave (CSW), which suffers from large optical transfer function (OTF) gap and poor spatial confinement^14,15^ (Figure S1F), and the Airy light sheet, whose center of mass shifts transversely with propagation (Supplementary Note 1; Figure S1G). Past works have attempted to overcome these limitations by coherently superposing multiple CSWs^18–21^. However, such coherent superpositions inevitably introduce intensity modulation along propagation that significantly shrinks the FOVs (Figure S2A).

As a result, most recent light sheet engineering efforts exploit additional spatial or temporal degrees of freedom. Notable examples include Bessel beam^3^ (Figure S2B), Airy beam^22^ (Figure S2C), lattice light sheet (LLS)^5^ (Figure S2D), Field Synthesis light sheet^9^ (Figure S2E), ‘space-time’ light sheet^17^, and multiphoton light sheet^4^. However, these approaches introduce spatially or temporally structured illumination that must be ‘smoothed out’ via beam scanning^3,5,9,22^ or the use of spectrally broadened light sources^4,17,23,24^. Consequently, the illumination duty cycle is reduced, peak intensity is increased, and photobleaching and phototoxicity are exacerbated^25–27^. Alternative approaches leveraging optical nonlinearities, such as soliton, were attempted to thwart diffractive spreading^28^, but these approaches often require nonlinear optical medium and are therefore incompatible with most biological specimen. Collectively, limitations of existing approaches reveal a fundamental and unsolved challenge in LSFM: achieving a light sheet that is simultaneously propagation-invariant, temporally invariant, and broadly compatible with diverse biological samples.

Another key limitation of LSFM is its susceptibility to optical aberrations. Refractive index (RI) inhomogeneities within multicellular organisms lead to aberrations that distort the light sheets and degrade resolution, contrast, signal-to-noise-ratio (SNR), and data interpretability with increasing imaging depth^11,29^. This degradation is particularly severe for nondiffracting light sheets that exploit the additional spatial or temporal degrees of freedom (Figure S2B-F). Integrating adaptive optics (AO) with LSFM offers a promising solution^29–31^. However, the associated instrumentation complexity, high cost, and reduced imaging speed—arising from the need to sequentially measure and correct spatially varying aberrations across image volumes—have limited its widespread adoption. There is an urgent but unmet need to develop light sheets that are intrinsically resilient to optical aberrations to enable rapid high-resolution intravital imaging.

Here, we introduced a fundamentally new class of light sheet—Polarization-Engineered Aberration-Resilient Light Sheet (PEARLS)—that addresses these long-standing challenges. By engineering the coherence structure of the optical field through polarization control, PEARLS generates a nondiffracting light sheet that is simultaneously propagation-invariant, temporally invariant, and intrinsically resilient to optical aberrations. Unlike existing nondiffracting light sheets, PEARLS requires neither beam scanning nor spectrally broadened light sources, enabling continuous illumination with full duty cycle and minimal photobleaching. We first verified key properties of PEARLS, including high resolution, spatiotemporally invariant profiles (Supplementary Note 1), minimal photobleaching, and intrinsic aberration resilience through both simulations and experiments. Then, we demonstrated the capability of PEARLS for noninvasive multicolor imaging of subcellular dynamics across diverse living systems, from cultured cells to intact organisms, spanning several orders of magnitude in space and time.

### Concept of PEARLS

The essence of PEARLS is that the incoherent superposition of multiple 1D CSWs with identical propagation length produces a light sheet whose profile remains invariant over the same propagation length. By engineering the properties of the constituent CSWs, we can tailor the light sheet profiles with high flexibility. For example, a CSW with high spatial resolution but poor confinement combined with another CSW that has lower resolution but stronger confinement can yield a light sheet with high resolution and strong spatial confinement (Figure 1A; Figures S2G,H & 3A-F).

**Figure 1.**
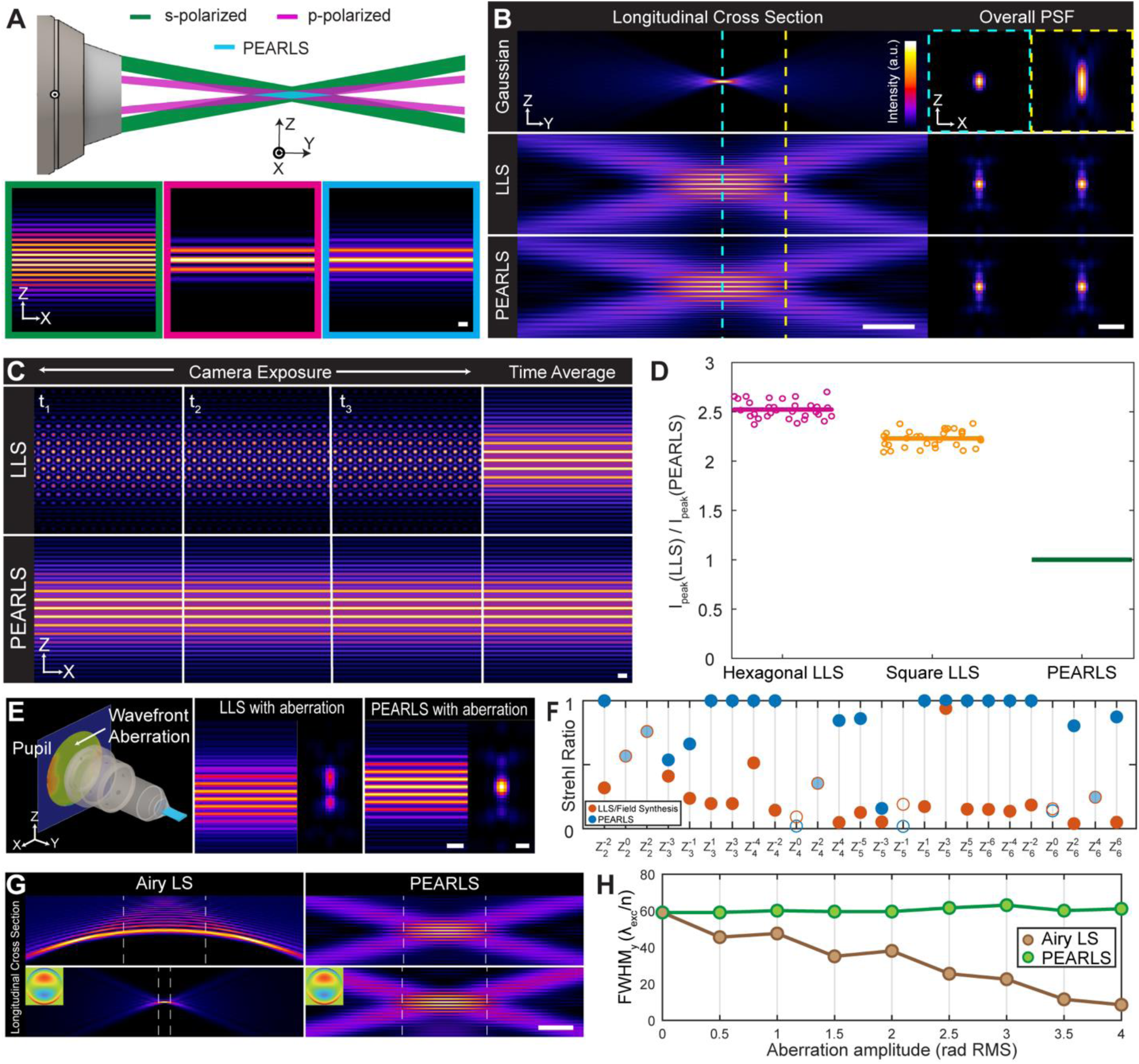
Polarization-Engineered Aberration-Resilient Light Sheet (PEARLS). (A) Principle of PEARLS: nondiffracting cosine standing wave (CSW) light sheets with orthogonal polarizations incoherently superpose. Green: s-polarized CSW, magenta: p-polarized CSW, blue: PEARLS. Scale bar: 2.5-λ_exc_/n. (B) Comparisons of light sheets with the same maximum numerical aperture (NA): Gaussian (top, NA=0.6), lattice light sheet (LLS, middle, NA_max_/NA_min_=0.6/0.56), and PEARLS (bottom, NA_max_/NA_min_=0.6/0.56). Left column shows the YZ cross-section, and right column the corresponding normalized overall XZ point spread functions (PSFs) at the foci (cyan dashed line) and at the half width at half maximum (HWHM) of the LLS/PEARLS (yellow dashed line) (Video S1). Scale bars: 25-λ_exc_/n (left) and 2.5-λ_exc_/n (right). (C) Instant and time averaged transverse (XZ) cross-sections of LLS and PEARLS. Scale bar: 2.5-λ_exc_/n. (D) Simulated peak illumination power ratios between LLS and corresponding PEARLS with the same spatial profiles and resulted image signal-to-noise-ratio (SNR). Each group contains various combinations of NA and Δ NA, ranging from 0.3-0.6 and 0.02-0.2, respectively. (E) Transverse cross-section profiles and overall PSFs of the LLS and PEARLS under an experimentally measured wavefront aberration. Scale bars: 5-λ_exc_/n (cross-section profiles) and 1-λ_exc_/n (overall PSF). (F) Strehl ratios of LLS/Field Synthesis and PEARLS under 25 different Zernike aberrations (Video S2). (G) Simulated YZ cross-sections of Airy light sheet (LS) and PEARLS without (top row) and with coma aberration (phase RMS=4 rad, bottom row). Scale bar: 25-λ_exc_/n. (H) Propagation length of Airy LS and PEARLS under coma aberrations with varying phase amplitudes. RMS: root-mean square. All results shown in this figure are based on numerical simulations.

This approach relies on two key physical conditions. First, all CSWs should have the same propagation length (Figure S3A-F). This is equivalent to requiring that the constituent CSWs possess the same momentum spreading along the propagation direction (Figure S3G-I). We derived the analytical expression governing this nondiffracting condition (Supplementary Note 2; Figure S3J,K). Second, the constituent CSWs need to be mutually incoherent. While temporal separation^9,32^, spectral separation^17,24^, and coherence control^23^ could in principle achieve this condition, they are suboptimal due to the need of spatiotemporal averaging or spectrally broadened light sources. Instead, we exploited a simple but robust physical property: optical fields with orthogonal polarizations are mutually incoherent (Figure 1A). Although this polarization-based approach limits the total number of simultaneously superposed CSWs to two, this is sufficient to generate any type of dithered LLS, including the recently invented harmonic balanced LLS^14^ (Supplementary Note 3; Figures S2G,H & 3L,M).

We first examined the propagation and temporal invariance of PEARLS. To benchmark its performance, we compared PEARLS with Gaussian light sheet—the most widely used LSFM—and LLS—the current performance standard. To ensure fair comparison and consistency with our experimental setups, we employed a numerical aperture (NA) of 0.6 on the illumination path and a NA of 1.0 on the detection path in our simulation. Unlike Gaussian light sheets whose axial resolution undergo a hyperbolic decline with propagation, PEARLS maintains its axial resolution over an extended propagation length (Figure 1B; Video S1). Furthermore, in contrast to LLS, which requires spatial averaging via galvo dithering^5^ (Figure 1C), or an alternative LLS realization named Field Synthesis, which requires temporal averaging via scanning in the Fourier space^9^ (Figure S4A), the transverse intensity profile of PEARLS is temporally invariant (Figure 1C), leading to full duty cycle illumination. As a result, to achieve the same average illumination power, the peak illumination power required for PEARLS is only 35%-50% of that required for LLS (Figure 1D) and 70%-80% of that required for Field Synthesis with identical cross-sectional profiles (Figure S4B).

### Aberration resilience and adaptive correction

Next, we systematically evaluated the aberration resilience of PEARLS. We hypothesized that two critical properties of PEARLS increase its tolerance to phase aberrations: (i) its reliance on incoherent superposition of CSWs, and (ii) all its wavevectors are confined in a 1D line in momentum space (k_x_=0). To test this hypothesis, we compared the performance of PEARLS and other light sheets under two types of aberrations: (1) experimentally measured aberrations from zebrafish, and (2) a comprehensive set of orthogonal Zernike aberration modes.

First, we introduced a wavefront aberration experimentally measured from a zebrafish spinal cord^33^ (Figure S5A) to a dithered LLS and PEARLS with identical profile (Figure 1E). Whereas the aberration significantly distorted both the light sheet profile and the overall point spread function (PSF) of LLS (Figure 1E, middle), PEARLS shows minimal changes in its profile and overall PSF (Figure 1E, right). To quantify how the resulted overall PSFs impact imaging performance, we carried out two analyses. First, we evaluated the axial resolvability of the LLS and PEARLS under this aberration using line patterns with variable spacings^14^. Whereas the resolvability of LLS deteriorated markedly under the aberration, PEARLS maintained nearly unchanged resolution (Figure S5C). Second, we simulated imaging of a field of sub-diffractive point emitters using the aberrated LLS and PEARLS with different SNRs (Figure S5D,F). Unlike LLS whose aberrated PSF led to misidentification of the emitters, PEARLS achieved results resembling the aberration-free case. To further analyze the origin of PEARLS’ aberration-resilience, we performed Fourier Shell Correlation analysis^34^. The analysis revealed that the middle k_z_ band of LLS generated by interference between the two side beams was strongly degraded by the aberration, whereas the corresponding spatial frequency components of PEARLS are almost unaffected (Figure S5E,G).

Second, we evaluated PEARLS’ resilience across the full space of common optical aberrations. Because any aberrations can be expressed as a linear combination of orthogonal and complete Zernike modes^31,35^, we performed a comprehensive mode-by-mode comparison across the first 28 Zernike aberrations (excluding piston, tip, and tilt). To quantify performance, we calculated the Strehl ratio—a standard measure of the optical system quality under aberrations^36^—of the resulting overall PSFs. For fair comparison, all nondiffracting light sheets were configured with identical NA (=0.6) and propagation length (=60λ_exc_/n; Figure 1F; Figure S6). Results highlighted several points. First, PEARLS is insensitive (Strehl ratio=1) to 11 of the 25 Zernike modes tested (Figure 1F). Second, PEARLS consistently outperforms all other light sheets evaluated. Compared with dithered LLS and Field Synthesis light sheet, PEARLS exhibited higher or equal Strehl ratios in 22 of the 25 Zernike modes (Figure 1F; Video S2). Similar results were obtained over scanning circular Gaussian beam light sheet (24 of 25 modes; Figure S6A), non-scanning Gaussian light sheet (24 of 25 modes; Figure S6B), double CSW light sheet (25 of 25 modes; Figure S6C), scanning Bessel beam light sheet (21 of 25 modes; Figure S6D), and single CSW light sheet (24 of 25 modes; Figure S6E). Last, these comparisons provide mechanistic insight into the origin of PEARLS’ aberration resilience. Notably, PEARLS outperforms double CSW light sheet, which shares the same electrical field intensity but lacks polarization separation, indicating the critical role of incoherent superposition. Furthermore, PEARLS outperforms dithered LLS and Field Synthesis light sheet with identical profile and OTF (Figure 1F), highlighting the importance of restricting all wavevectors to a 1D line (k_x_=0).

Third, we investigated how aberrations affect the propagation-invariant property of PEARLS. Unlike Airy light sheet whose propagation-invariance relies on a precisely defined cubic pupil phase, the propagation-invariant property of PEARLS arises because all constituent beamlets share the same longitudinal momentum spread (Δk_y_). Consequently, aberrations are expected to have minimal impact on its propagation length. Consistent with this expectation, the propagation length of PEARLS remained largely unchanged across all 25 Zernike modes (Figure S7A,B). In contrast, the propagation length of Airy light sheet changed substantially under several aberration modes (Figure S7C,D). In particular, severe coma aberration can completely disrupt the nondiffracting propagation of Airy light sheet, effectively transforming it to a Gaussian-like light sheet with over six-fold reduction in propagation length (Figure 1G,H). By comparison, PEARLS showed minimal propagation length change under the same coma aberration (Figure 1G,H).

Last, we explored simple strategies to adaptively correct residual aberration effects in PEARLS. Because PEARLS consists of two incoherently superposed CSWs, we hypothesized that aberrations primarily manifest as two kinds of relative misalignments between the constituent CSWs: (i) a relative defocus along the propagation direction (Y-axis), and (ii) a relative axial shift (Z-axis) (Figure S8F). These effects can be compensated by translating one CSW relative to the other along Y- and Z-axes (Figure S8F) without the need for complex hardware such as spatial light modulator (SLM) or deformable mirror. To test this hypothesis, we performed numerical simulations and confirmed that this translation-based correction is able to substantially restore PEARLS’ profile, achieving Strehl ratios greater than 0.9 across all 25 Zernike modes tested (Figure S8G) with residual deviations arising from aberrations within each beamlet.

### Experimental implementation and characterization

We experimentally implemented PEARLS using only off-the-shelf optics, avoiding programmable element or synchronized galvo scanning for broad adoption (Figure 2A; Figure S8A). An overview of the system highlights several key features. First, to attain mutual incoherence, we generated the two CSWs using a pair of independently translatable Fresnel biprisms (Figure 2A, FBP1&2) that are separated by a polarizing beam splitter (Figure 2A, PBS3). Second, we achieved the nondiffracting condition with a variable beam expander (Figure 2A, VBE2). Third, unlike LLS whose width is constrained by the number of pixels of an SLM, PEARLS generation does not rely on active element and therefore its width is, in principle, unconstrained. Here, we demonstrated a 750-μm wide PEARLS with uniform intensity profile by pairing a Powell lens with a cylindrical lens (Figure 2A, PL&CL1; Figure S8B,C). Last and most importantly, the absence of a photomask and diffractive components leads to minimal light loss with a transmission efficiency of ∼85%. For comparison, the beam shaping units of LLS and Field Synthesis have a transmission efficiency of ∼3% and ∼35%, respectively^9^.

**Figure 2.**
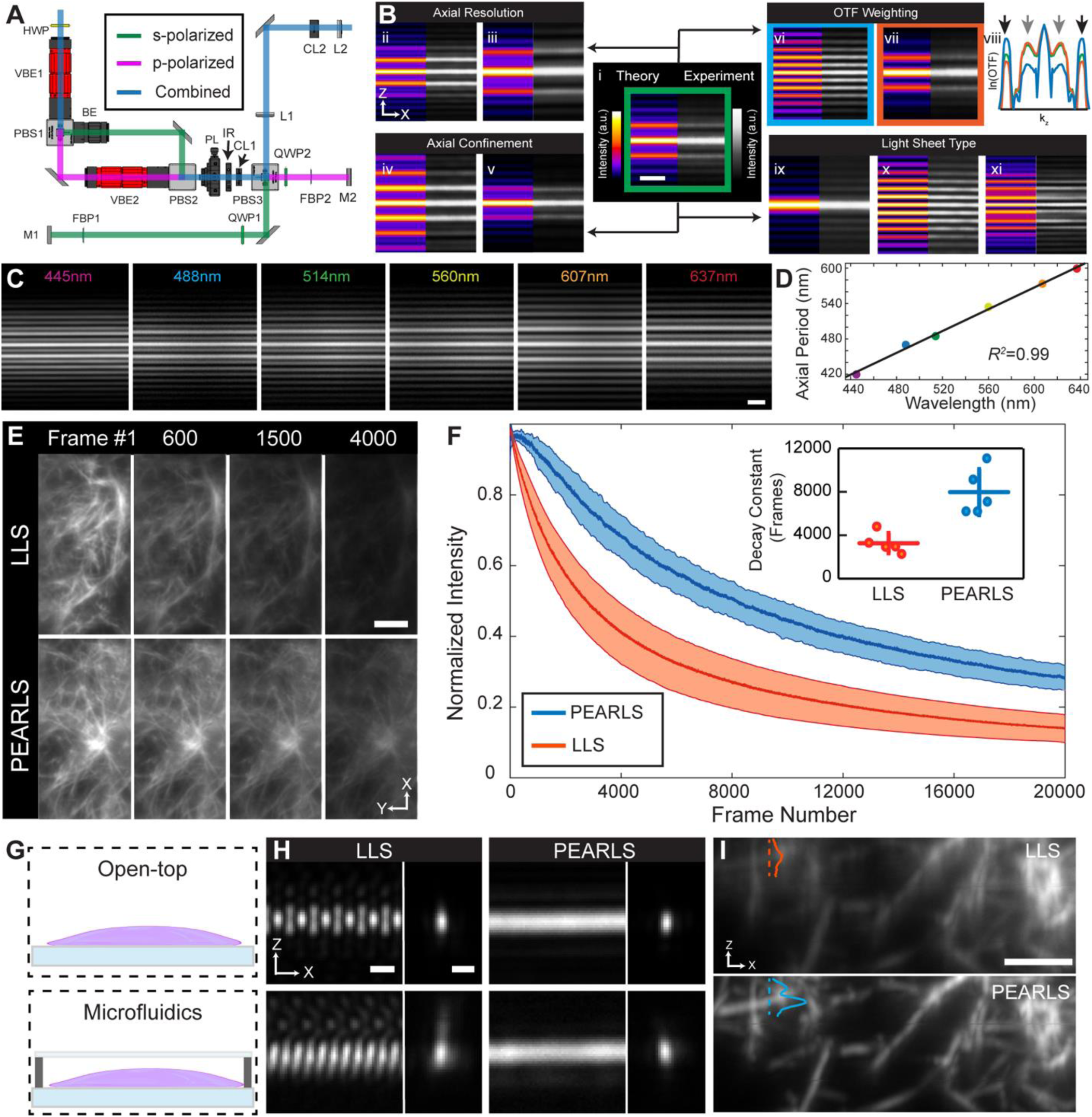
Experimental characterizations. (A) Optical layout of the beam shaping unit of PEARLS. HWP, half-wave plate; VBE, variable beam expander; PBS, polarizing beamsplitter; BE, beam expander; PL, Powell lens; IR, iris; CL, cylindrical lens; L, lens; QWP, quarter-wave plate; FBP, Fresnel biprism; M, mirror. (B) Demonstrations of PEARLS’ tunability of axial resolution (i-iii, NA_max_/NA_min_=0.46/0.38, 0.6/0.52, and 0.36/0.28); confinement (iv-v, NA_max_/NA_min_=0.46/0.42 and 0.5/0.38); optical transfer function (OTF) weighting (NA_max_/NA_min_=0.46/0.38, intensity ratio of outer to inner CSW on the rear pupil plane is 1:1, 4:1, 1:2 for green (i), blue (vi), and orange (vii), respectively). Arrows in (viii) indicate the axial spatial frequency contributions from the individual CSW components (black: outer CSW; gray: inner CSW); and light sheet types (i, hexagonal-type, ix, square-type, x, CSW, and xi, a profile without an equivalent LLS). Simulations (left) and experiments (right) show good agreement. Scale bar: 5-λ_exc_ /n. (C) PEARLS light sheet profiles (hexagonal type, NA_max_/NA_min_=0.6/0.5) at six different wavelengths, ranging from 445, 488, 514, 560, 607, and 637nm. Scale bar: 2-μm. (D) Relationship between the wavelength and the corresponding axial period. Linear fitting yields *R*² = 0.99. (E) Photobleaching comparisons of hexagonal type PEARLS and dithered LLS with identical profile (Video S3). Each frame was normalized with respect to (w.r.t.) the maximum intensity of the Frame 1 of each light sheet. Representative Frames 1, 600, 1500, and 4000 taken by LLS (top) and PEARLS (bottom) were shown. Scale bar: 5-μm. (F) Averaged photobleaching curves of PEARLS (blue) and LLS (red). The shaded areas indicate ±1 standard deviation (SD) from five datasets taken at five different regions of the same collagen gel with intensities normalized. Inset: characteristic decay constants of LLS (red) and PEARLS (blue) obtained by fitting individual photobleaching curves with exponential decays. Horizontal and vertical lines indicate the mean and ±1 SD of the five datasets, respectively. (G) Comparison of PEARLS and LLS applied to imaging within a microfluidic device. (H) Transverse cross-section profiles and overall XZ PSFs of LLS (left) and PEARLS (right) without (top) and with (bottom) the microfluidic device. Scale bars: 1-μm. (I) XZ views of raw collagen images within the microfluidic device taken by LLS (top) and PEARLS (bottom) with intensity normalized w.r.t. the maximum intensities of each volume. Scale bar: 5-μm. Red and blue curves show intensities of the corresponding line cuts. Raw collagen images are shown and used for analysis in (E, F, I).

Despite the absence of programmable elements, PEARLS is capable of fully control the light sheet profiles, including axial resolution (Figure 2B, i-iii), axial confinement (Figure 2B, i, iv, v), relative weighting of different spatial frequency components (Figure 2B, i, vi-vii), and light sheet types (Figure 2B, i, ix-xi). Notably, because of the conceptual simplicity and compact implementation of PEARLS, experimentally measured beam profiles closely match the simulated profiles, both at the focal plane and half-width half maximum (HWHM) along the propagation direction (Figure 2B; Figure S8D,E). Furthermore, PEARLS uses only refractive, not diffractive, elements, enabling simultaneous multi-color illuminations. We experimentally characterized six co-aligned PEARLSs spanning wavelengths from 445-nm to 637-nm (Figure 2C). Quantitative measurements show that the axial periods scale linearly with wavelength (*R*^2^=0.99; Figure 2D), confirming minimal chromatic aberrations in the system.

Last, our implementation of PEARLS enabled straightforward correction of optical aberrations. As a demonstration, we introduced aberrations by inserting a transparent plastic film with uneven surface. By adjusting VBE2 and a mirror (Figure 2A, M2) conjugated to the rear pupil, we translated the p-polarized CSW along the propagation and axial directions, respectively. These simple adjustments largely restored the beam profile, yielding a cross-correlation coefficient (Pearson’s r) of 0.92 relative to the aberration-free case (Figure S8H,I).

### Comparisons with LLS microscopy in photobleaching

Because LLS microscopy remains a widely used technology for 3D subcellular dynamics imaging, we compared PEARLS and LLS in imaging samples containing sub-diffractive features (Figure S9A). For fair comparison, we integrated our PEARLS beam shaping unit (Figure S8A) into an LLS microscope so that PEARLS and LLS share the same optical path except for the beam shaping units. PEARLS and dithered LLS with identical profiles and therefore identical overall PSFs (Figure S9B,C; Table S1) were applied to image Alexa Fluor 488 labeled collagen gels whose structures mimic the extracellular matrix and offer a reproducible imaging standard^37^. 3D image stacks and orthogonal slices confirmed that the two light sheets produced similar spatial resolution in all three dimensions (Figure S9D-G).

Next, we focus on quantifying the photobleaching of PEARLS and LLS. Our simulation studies showed that, to achieve the same SNR, the required peak illumination power of PEARLS is only 35%-50% of that of LLS (Figure 1D). Because high peak illumination power exacerbates photobleaching^25–27^, we hypothesized that PEARLS would exhibit reduced photobleaching. To test this hypothesis, we performed rapid time-lapse imaging of collagen gels using PEARLS and dithered hexagonal LLS with identical profile (Figure 2E; Video S3). To minimize the influence of sample heterogeneity, we applied each light sheet at five different regions of the gel and extracted corresponding photobleaching curves to determine characteristic decay constants (Figure 2F; Methods). In comparison to LLS, PEARLS exhibits statistically significant reduced photobleaching with ∼2.5-fold larger characteristic decay constant (Figure 2F, inset). We obtained similar results when comparing PEARLS with dithered LLS with identical profile (Figure S9H; Video S4). In addition to reducing photobleaching within the imaging FOV (i.e., the camera’s acquisition region), PEARLS eliminates unnecessary lateral (X-axis) photobleaching outside the imaging FOV (Figure S9I). Such out-of-FOV photobleaching is particularly problematic when spatial tiling is used to image a large volume^11^. In LLS, beamlets with non-zero lateral momentum inevitably illuminate regions outside the imaging FOV (Figure S9I, left panel). In contrast, because all constituent CSWs of PEARLS have zero lateral momentum (k_x_=0), illumination of PEARLS is strictly confined within the imaging FOV (Figure S9I, right panel). Systematic analysis of LLS with different lengths and widths showed that: narrow LLS, which is used for imaging multicellular organisms due to the limited size of isoplanatic patch^29^, and long LLS, which is used for cleared or expanded tissues^14,38^, can have out-of-FOV illumination area comparable to the imaging FOV. PEARLS, by contrast, produces no lateral photobleaching outside the imaging FOV regardless of the beam profiles (Figure S9J). In short, in comparison to LLS, PEARLS exhibits both reduced photobleaching within the imaging FOV and no lateral photobleaching beyond the imaging FOV.

### Imaging through microfluidic device, changing medium, and inverted configuration

Microfluidics devices have attracted growing attentions in biomedical research because of its low sample consumption, high efficiency, and multifunction integrability^39^. However, LSFM imaging through microfluidic devices is often compromised by optical aberrations arising from the RI mismatch between microfluidics devices and surrounding medium^40^. Given the aberration-resilience of PEARLS, we hypothesized that it would enable high resolution imaging inside microfluidic devices with minimal correction.

To test this hypothesis, we created a microfluidic device incorporating a ∼50-µm-thick fluorinated ethylene propylene (FEP) cover^40^, and compared the performance of PEARLS with dithered LLS by imaging fluorescently labeled beads and collagen gels inside the device (Figure 2G). Because PEARLS improves the robustness of the excitation light sheet while leaving detection-path aberrations unchanged, we corrected detection-side aberrations for both PEARLS and LLS using phase retrieval^41^ and a deformable mirror. Minor aberrations on the detection path remained likely due to attenuation of marginal rays^40^. Measurement of the light sheets profiles and overall PSFs showed that the microfluidic devices distorted the LLS, whereas PEARLS maintained a largely unchanged beam profile and overall PSF (Figure 2H). Consequently, collagen gel imaged with LLS exhibited degraded structural definition, while PEARLS was noticeably less affected and preserved fine structural details (Figure 2I).

Last, we experimentally demonstrated the aberration correction capability of PEARLS under two commonly encountered imaging scenarios: exchange of immersion media and imaging through coverslip. First, when the immersion medium was switched from 1× phosphate-buffered saline (PBS) to deionized water (Figure S8J), the RI mismatch introduced spherical aberration (𝑍^0^), a term known to substantially affect PEARLS’ profile (Figure 1F). This aberration induced a relative shift between the two CSWs along the propagation axis (Figure S8K, red box). A simple tuning of VBE2 restored the desired beam profile (Figure S8K, green box), yielding a cross-correlation coefficient of 0.98 relative to the 1×PBS reference (Figure S8L). Second, we examined correction through a standard 170-µm glass coverslip (Figure S8M,N). The resulting aberration is more complex due to the oblique geometry between the objectives and the coverslip^42–44^. To correct, we adopted a two-step correction procedure: first adjusting global beam translations followed by adjusting the relative shifts between the two CSWs. Despite some residual distortions, this procedure can largely restore the beam profile (Figure S8N), improving the cross-correlation coefficient from 0.80 to 0.95 relative to the upright configuration reference (Figure S8O). Together, these experiments demonstrate that the intrinsic aberration-resilience property and simple correction strategy of PEARLS can satisfy imaging needs across diverse experimental configurations, including different types of sample holders, immersion medium, and objective configurations.

### Observation of rapid subcellular dynamics in cultured cells

The high axial resolution, spatiotemporally invariant profile, and aberration-resilience of PEARLS could enable prolonged noninvasive observations of living cells within microfluidic devices. As a demonstration, we imaged HeLa cells stably expressing mCherry-CAAX for 300 image volumes (251 images per volume) at 13-s intervals (Figure 3A; Figure S10; Video S5). Orthoslice with sub-diffractive features showed axial resolution (Figure 3B). No detectable photobleaching or perturbation on cellular dynamics were observed. Intercellular interactions mediated by rapid filopodia movements were observed at sub-minute timesscales (Figure 3C), highlighting the dynamic and interactive nature of HeLa cells.

**Figure 3.**
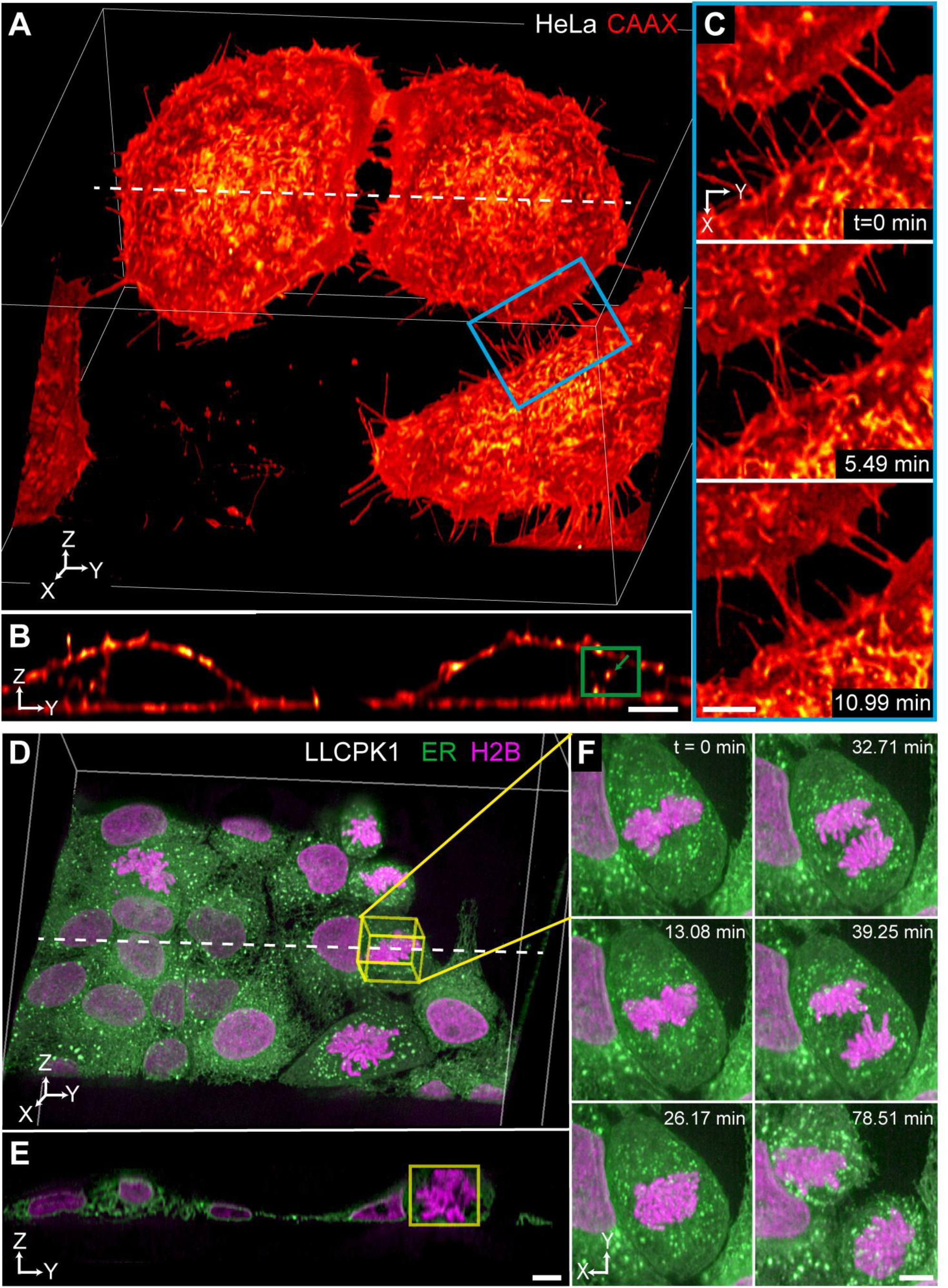
Three-dimension intercellular and intracellular dynamics. (A) Volume rendering of live HeLa cells transfected with mCherry-CAAX showing rapid membrane dynamics and intercellular filopodia interactions. This is the first time point from a time-lapse data with 300 time points (Video S5). Bounding box: 111-μm by 110-μm by 100-μm. (B) YZ orthogonal view of white dashed line in (A), showing clearly resolved cell membranes. A diffraction-limited feature (green arrow) highlights 3D resolution of PEARLS. Scale bar: 5-μm. (C) Zoom-in time-lapse images of the blue box in (A), showing intercellular interactions via filopodia. Scale bar: 5-μm. (D) Simultaneous two-color volume rendering of LLCPK1 cells expressing mEmerald-ER (green) and mCherry-H2B (magenta) undergoing cell division. This is the first time point from a time-lapse data with 400 time points (Video S6). Bounding box: 111- μm by 110-μm by 100-μm. (E) YZ orthogonal of white dashed line in (D), highlighting a mitotic cell (yellow box). Scale bar: 5.0-μm. (F) Zoom-in time-lapse images of the mitotic cell in (D and E), covering metaphase to cytokinesis. Scale bar: 20-μm. All images shown were deconvolved.

Mitosis is a spatially complex, temporally dynamic, and phototoxicity sensitive process that is central to cellular functions. A key advantage of fluorescent imaging is that the spectral diversity of fluorescent proteins enables observations of the interactions between subcellular components. We performed simultaneous two-color imaging of LLC-PK1 cells that are stably transfected with mEmerald-ER and mCherry-H2B with a total of 400 volumes (251 images per volume) at 16-s interval over 2 hours (Figure 3D; Video S6). Mitotic cells with clear cell rounding and chromosome condensing were identified (Figure 3E). We tracked the mitosis of a dividing from metaphase all the way to cytokinesis (Figure 3F) and observed complete exclusion of endoplasmic reticulum (ER) by the chromosomes during anaphase and telophase.

### Noninvasive long-term tracking of embryonic developments

While cultured cells have offered critical biological insights, cells inside our bodies do not live in isolation and are in constant interactions with neighboring cells and extracellular matrix. Therefore, it remains desired to directly observed cellular processes within the native multicellular environments, where all the cues that drive phenotypes are present^29,45^. Here, we applied PEARLS for imaging of subcellular dynamics in two widely used model organisms: zebrafish and *Drosophila melanogaster*. We focused on two critical cellular processes—mitosis and migration—in early embryonic developments. No aberration correction was applied to the light sheet illumination path, and only detection-path corrections were performed using a deformable mirror.

Zebrafish (*Danio Rerio*), a transparent vertebrate that shares a high degree of genomic and biochemical similarities with human, represent an ideal model organism to study development^46^. We performed 3D time-lapse imaging of pan-nuclear, GFP-histone labeled transgenic zebrafish embryos undergoing late discoidal meroblastic cleavage and mid-blastula transition. First, we directly compared the *in vivo* aberration-resilience of PEARLS and dithered LLS with identical profiles at the same imaging positions (Figure S11A). No excitation path correction was applied to either PEARLS or LLS, and the same deformable mirror correction was applied to the shared detection path. Whereas PEARLS can clearly resolve chromosomal structures, LLS exhibited noticeable axial blurring and reduced structural definition (Figure S11B).

Next, we recorded an imaging volume of 220-µm × 250-µm × 30-µm (626 images per volume) for ∼2.5-hour at 26-s interval with a total of 322 volumes (Figure 4A; Video S7). Multiple diffraction-limited spots could be seen from the 3D visualization, confirming PEARLS’ high axial resolution and aberration-resilience (Figure S11C,D). 3D chromosomal dynamics, including condensation, alignment, and separation of sister chromatids, from dozens of mitotic cells were clearly visualized and tracked (Figure 4B). At the onset of the session, cells within the blastula divide meta-synchronously, characteristic of late cleavage and early blastula periods. This synchrony gradually lost with lengthening of cell cycles in later imaging session, marking the onset of mid-blastula transition. Over the entire imaging session, we observed less than 10% change in the averaged fluorescence intensity, indicating minimal photobleaching (Figure S11E).

**Figure 4.**
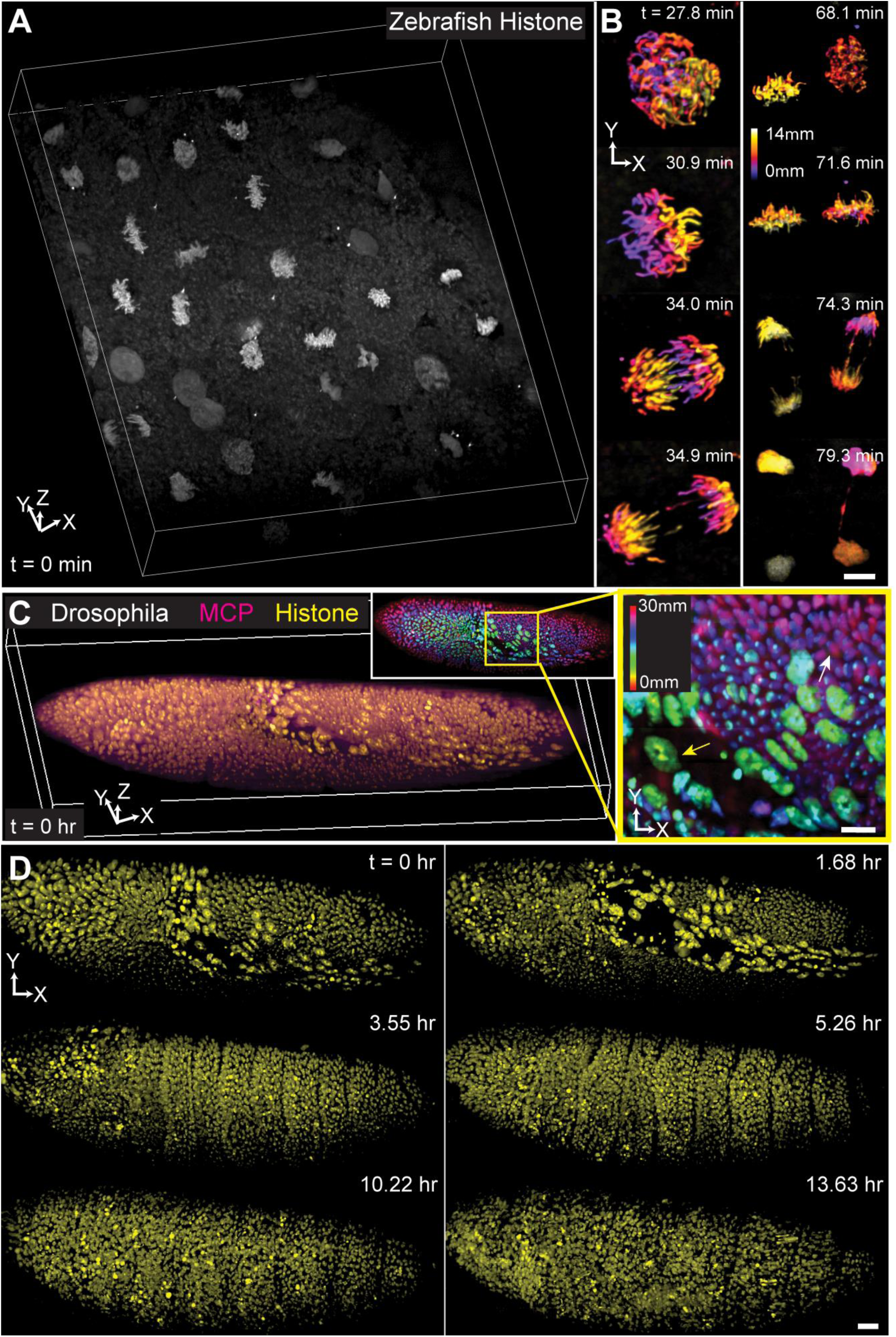
Cell divisions and migrations in embryogenesis. (A) Volume rendering of a pan-nuclear (H2B-GFP) zebrafish embryo (∼3hpf), highlighting synchronized mitosis. This is the first time point from a time-lapse data with 322 time points (Video S7). Bounding box: 264-μm by 221-μm by 60-μm. (B) Selected frames showing mitotic progression—chromosomal condensation, alignment, and separation— in two representative regions. Left: a single dividing cell; right: two adjacent cells undergoing synchronized division. Colors encode depth along the Z-axis. Scale bar: 5-μm. (C) Volume rendering of simultaneous two-color imaging of *Drosophila melanogaster* embryo expressing mNeonGreen-MCP (magenta) and mRFP-histone (yellow), imaged with square-type PEARLS (NAmax/NAmin=0.42/0.38). This is the first time point from a time-lapse of 412 time points (Video S8). Bounding box: 432-μm by 194-μm by 60-μm. Inset: maximum intensity projection of the XY view. Colors encode depth along the Z-axis. A zoom-in of the yellow box is shown on the right, highlighting drastic nucleus size difference between cells in amnioserosa (yellow arrow) and cells in germ band (white arrow). Scale bar: 10-μm. (D) Lateral views of selected frames showing germ band extension (first row), formation of body segments (second row), and coordinated muscle contractions (third row). All images shown were deconvolved.

Finally, we performed simultaneous two-color, long-term imaging of *Drosophila* embryogenesis. *Drosophila* is a widely used model organism thanks to its genetic accessibility and rapid development. Transgenic *Drosophila* embryos expressing mNeonGreen-MS2 coat protein (MCP) and mRFP-histone^47^ were dechorionated and mounted on a coverslip. We performed two-color imaging of a *Drosophila* embryo undergoing gastrulation and subsequent morphogenesis over a 220-µm × 450-µm × 30-µm volume with 1126 images per volume for ∼14-hour at 2-min interval with a total of 412 imaging volumes (Figure 4C; Video S8). Early in the imaging session, amnioserosa, an extraembryonic tissue composed of large cells (yellow arrow, Figure 4C), underwent dynamic rearrangements while the germ band, a collection of smaller cells (white arrow, Figure 4C), began to extend along the embryo’s ventral surface (first row, Figure 4D). Afterwards, the gradual formation of body segments became clearly visible as repeated bands of nuclei in the ectoderm, reflecting early patterning events in the embryo (second row, Figure 4D). Last, coordinated muscle contractions from the activation of the neuromuscular system led to rapid twitching of the entire embryo (third row, Figure 4D). Throughout the 14-hour imaging session, the image quality remained stable (Figure S11F), despite complex, time-varying aberrations, illustrating the aberration resilience of PEARLS.

## Discussion

PEARLS is enabled by two key conceptual advances. First, it generates a spatiotemporally invariant profile through the incoherent superposition of orthogonally polarized CSWs that satisfy the nondiffracting condition. This spatiotemporal invariant profile grants PEARLS a combination of high spatial resolution, large FOV, and low photobleaching and phototoxicity. At equivalent resolution, PEARLS offers 5-10-fold larger FOV than Gaussian light sheets, thus 5-10-fold improvements in acquisition speed. Compared with LLS of identical profile, PEARLS can achieve the same SNR with only 35-50% of the peak illumination power, resulting in ∼2.5-fold slower photobleaching. Importantly, the full-duty-cycle illumination of PEARLS places it near the theoretical lower bound of photobleaching achievable in light-sheet with a given profile. Second, the incoherent superposition of CSWs with zero lateral momentum makes PEARLS intrinsically more resilient against optical aberration in comparison to all other light sheets. Across a comprehensive set of Zernike aberration modes, PEARLS exhibited better or equal performance in 22 of 25 modes compared to LLS and Field Synthesis, in 24 of 25 modes compared to Gaussian light sheet, 25 of 25 modes compared to double-CSW light sheet, and 21 of 25 modes compared to Bessel beam light sheet, and 24 of 25 modes compared to single-CSW light sheet (Figure S12A). This aberration-resilience enabled high resolution imaging in optically challenging environments such as microfluidics devices and multicellular organisms. In addition, PEARLS spatially patterned coherence strategy enables the generation of lateral interference patterns for structured illumination based super-resolution imaging, a capability shared with LLS but not achievable by Field Synthesis (Figure S12B).

In addition to the conceptual advances, PEARLS offers several practical advantages. First, the absence of photomasks and diffractive components yields high transmission efficiency (∼85%), far exceeding that of LLS (∼3%) and Field Synthesis (∼35%) (Figure S12C). Second, unlike LLS whose light sheet width is limited by SLM pixel count, PEARLS’ width is in principle unconstrained. Third, PEARLS uses only off- the-shelf refractive optics, permitting simultaneous multicolor imaging and allows simple correction of optical aberrations, including immersion medium exchange and imaging through coverslips. Last but importantly, PEARLS is readily accessible to a broad user base. Unlike LLS and Field Synthesis systems that require complex optical setups and instrument synchronization, PEARLS can be implemented as a simple modification of a conventional Gaussian light sheet microscope, requiring only a compact optical module at the rear pupil plane (Figure S12D). This ease of conversion can enhance its impact and accessibility to the broad LS community.

We note several limitations of PEARLS and opportunities for further development. First, PEARLS utilizes polarization to achieve mutual incoherence, constraining its constituent CSWs to two. Future work could explore additional degrees of freedom of light to generate more complex light sheet profiles^48,49^. Second, while PEARLS is more aberration-resilient than all existing light sheets, it can still suffer from performance degradation in highly scattering samples^50–52^. Integration with near infrared multiphoton lasers is a natural step forward^4,53,54^. PEARLS’ high transmission efficiency with limited dispersion makes multiphoton PEARLS feasible without expensive high-power lasers. Last, like every LSFM, the huge amount of data generated by PEARLS poses a pressing challenge for image processing, visualization, and analysis, calling for versatile software pipelines^55–57^. Despite these future possibilities, the demonstrated advantages of PEARLS already open up opportunities for both basic biological research and biomedical studies.

## Methods

### Numerical simulations of light sheet profiles

We simulated the illumination profiles of different light sheets at the sample by first defining the complex electrical fields at the rear pupil plane of the illumination objective (Figure S1C). For beams or beamlets with top-hat profiles, we used a constant electrical field amplitude within the NA broadening of each beam or beamlet. For beams or beamlets with Gaussian profiles, we used a Gaussian electrical field amplitude with its amplitude dropping to 1/e within the NA broadening of each beam or beamlet. We calculated the electric field on the sample plane by Fourier transforming the electrical field at the rear pupil plane. We then calculated the intensity profile by taking the modulus square of the sample plane electrical field. To simulate beam profiles at different locations along propagation at the sample space, we followed the angular spectrum method by multiplying a defocus factor exp(2π*ik*_y_*y*) to the rear pupil electrical field prior to the Fourier transformation^16,58^. We simulated the detection PSF based on the annular field integrals formulated by Richards and Wolf^59^ with an NA of 1.0 and a RI of 1.33. The overall PSF is the product of the illumination profile and the detection PSF. The OTF is the Fourier transform of the overall PSF. The light sheet propagation length is defined as twice the Y-axis distance from the focal plane (Y=0) to where the intensity falls to half (Y_HWHM_). To quantify light sheet confinement, we computed the cumulative energy of the overall PSF by integrating the intensity along the Z-axis and calculate the range in which half of the energy is distributed (I_FWHM_).

### Aberration simulation, Strehl ratios and spatial resolvability

To numerically analyze the effect of optical aberration on light sheet, we added a complex phase factor described by the first 28 Zernike polynomials to the rear pupil electrical field. The Zernike phases were generated by a modified MATLAB *zernfun* function, where each mode was normalized such that its root mean square (rms) equals 1 within the unit disk, which is defined by the NA_max_ of the beam. We then multiplied each normalized Zernike mode by a constant of 6 to obtain larger aberrations. With these added Zernike phases, we simulated the aberrated overall PSFs and quantified the degree of optical aberrations using Strehl ratio. We adopted the Strehl ratio definition as the ratio of the peak intensity of the overall PSF on the focal plane of the illumination objective (y=0) with and without aberration. In addition to the Strehl ratio, we numerically simulated a resolution target by convoluting the self-normalized overall PSF with a line pattern of variable spacing (line spacing=110-nm, 220-nm, 360-nm, 530-nm, 730-nm, 960-nm, 1.22-µm, 1.51-µm, 1.83-µm, 2.18-µm, 2.56-µm, 2.97-µm) as described in Liu et al.^14^. To better simulate realistic imaging scenario, we added Poisson noise with a SNR of 3.

### Simulated fluorescent beads images and Fourier shell correlation

To determine the degree of discrepancy with and without optical aberration, we simulated 3D volumes of fluorescent beads and applied the simulated images to Fourier shell correlation (FSC) algorithm. We began the simulation of fluorescent beads by defining an image of the ground truth: a randomly distributed pixels with a certain preselected density. We then convolved the ground truth image with the 3D unaberrated or aberrated overall PSFs. Afterwards, we added Poisson noises to the image^16^. To start FSC, we first generated two simulated fluorescent beads, ***I***_1_(x, y, z) and ***I***_2_(x, y, z), volumes with the same SNR but with independent ground truth along with their Fourier transforms, denoted as ***F***_1_(k_x_, k_y_, k_z_) and ***F***_2_(k_x_, k_y_, k_z_). We then followed the FSC measure developed by Koho et al.^60^. Because we are mostly interested in understanding how optical aberrations affect the axial resolution, we averaged the results azimuthally as well asl between the positive and negative k_z_ to generate a map of Fourier correlation, FC (k_r_, │k_z_│). In addition, we applied a mask based on the thresholding of the ideal unaberrated OTF to eliminate the interference of noise by setting the boundary of the resolvable resolution on the FC map.

### Microscope hardware

A laser combiner supplies the microscope with visible excitation. Linearly polarized continuous-wave lasers at 405nm, 100mW (TOPTICA, IBEAM-SMART-405-S-BZ), 445nm, 100mW (IBEAM-SMART-445-S-BZ), 488nm, 500mW (MPB, 2RU-VFL-P-500-488-B1R), 514nm, 1000mW (MPB, 2RU-VFL-P-1000-514-B1R), 560nm, 1000mW (MPB, 2RU-VFL-P-1000-560-B1R), 607nm, 1000mW (MPB, 2RU-VFL-P-1000-607-B1R), 642nm, 2000mW (MPB, 2RU-VFL-P-2000-6224-B1R). Linearly polarized lasers with a 1.12 mm FWHM diameter exit the laser combiner are coaligned and expanded by 2×. Each laser is free space coupled and independently aligned to a single overlapping beam path. The intensity of each laser entering the microscope is independently controlled and fast modulated using an Acoustic Optical Tunable Filter (AOTF; AA, AOTFnC-400.650-CPCh-TN) driven by a 4 channel Multi-Purpose Digital Synthesizer (AA, MPDS4C-B66-22-52.111). The laser beams then enter either the PEARLS beam shaping unit (described below) or the LLS beam shaping unit (described below). The two beam shaping units merge at a custom photomask conjugated to the rear pupil plane: for PEARLS, the photomask is set at an empty position. For LLS, annular mask patterns with different NA_max_ and NA_min_ were used. A transform lens (FL=150-mm) routes from the photomask to a sample-conjugate 4kHz resonant galvo (Cambridge Technology, CRS series, 6SC04KA040-01Y) that wobbles the light sheets to reduce striping artifacts from absorption and scattering within the multicellular organisms. A transform lens (EFL=80-mm) routes the lasers to a pupil-conjugated Z-scan galvo (Cambridge Technology, 6215H series, 6SD11226) that moves the light sheet axially. The laser beams are then imaged via 4F lens relay (EFL=150-mm, FL=150-mm) to a pupil-conjugated X-scan galvo (Cambridge Technology, 6215H series, 6SD11587) that dithers the LLS to smooth out the intensity alone the light sheet width direction. For PEARLS, the X-scan galvo was kept steady. Another 4F lens relay (EFL=150-mm, EFL=400-mm) relays the light to the pupil of the excitation objective (Thorlabs, 20X, 0.6NA). Emitted fluorescence is captured by the detection objective (Zeiss 421452-9800-000, 20×, 1.0NA) and is relayed by the tube lens (Zeiss 423731-8246-000, EFL=165mm) and a pair of closely placed lenses (ELF=100-mm) to a deformable mirror (ALPAO, DM69). A custom long-pass dichroic mirror (Semrock, DI03-R561-T3-32×40) separates the emission light into two different camera paths. The light of each path is focused by a long focal length lens (EFL=300-mm) into an sCMOS camera (Hamamatsu ORCA-Flash4.0 V3).

### PEARLS beam shaping unit

Co-aligned lasers exiting the laser combiner and AOTF passes through an achromatic half-wave plate (Bolder Vision Optik AHWP3) that controls the relative weighting of the two CSWs. We designed the system that can break down into three different modules: (i) a beam size control module, (ii) a light sheet generation module, and (iii) a CSWs generation module. First, the beam passes through a beam size module that defines the confinement of the light sheet: an achromatic variable beam expander (0.5×-2.5× Thorlabs, ZBE1A) that controls the input beam size by adjusting the magnifying collar. Then the beam is separated by polarization via a polarizing beamsplitter cube (Thorlabs, PBS251) to get their beam sizes tuned separately. The beam on path one (s-polarized, Figure 2A) passes through an achromatic fixed magnification beam expander (2×, Thorlabs GBE02-A) inversely to get demagnified (0.5×). The beam on the second path (p-polarized, Figure 2A) passes through an achromatic variable beam expander (Thorlabs, ZBE1A). Last, the two beams are recombined by a second polarizing beamsplitter cube (Thorlabs, PBS251). The second module creates the light sheet: the recombined beam then passes through a Powell lens (20-degree fan angle, Laserline Optics LOCP-8.9R20-2.0) that expands the beam into a uniform line illumination on the x-direction, followed by an iris that controls the light sheet width, and an achromatic cylindrical lens (EFL=50 mm) that collimates the beam on the x-direction. Lastly, the light sheet beam enters the third module to generate the CSWs: a third polarizing beamsplitter cube (Thorlabs, PBS251) splits the beam again into orthogonal polarization paths—each path is composed of an achromatic quarter-wave plate (Bolder Vision Optik AHWP3)—to form an optical insulator, a Fresnel biprism (170-degree, Newlight Photonics FBP2020G-170) with custom antireflective coating that splits the light in z-direction, and a mirror. The two beams are recombined a second time by the third polarizing beamsplitter cube, passes through a pair of 4F lens (EFL=200-mm) with a cylindrical lens in between (EFL=75mm). The 4F lens pairs relay the mirror of the p-polarized path onto the pupil conjugated photomask. The cylindrical lens in between forms a 4F relay in the x-direction with the first lens.

### LLS beam shaping unit

Co-aligned lasers exiting the laser combiner and AOTF are shaped by passing through a Powell lens (20-degree fan angle, Laserline Optics LOCP-8.9R20-2.0) that expands the beam into a uniform line illumination on the x-direction followed by a spherical lens (FL=50-mm) that collimates the beam on the x-direction and focuses the beam on z-direction. A pair of cylindrical lenses (FL=50-mm and FL=300-mm) then focuses the light on a sample-conjugated spatial light modulator (SLM, Meadowlark Optics, P1920-0635-HDMI, 1920 × 1152 pixels). Before the lasers entering the SLM, they pass through a custom dichroic stack composed of a series of long-pass or band-pass dichroics and mirror (Di03-R405/Di03-R442/Di03-R488/Di03-R532/Di03-R561/Di01-R488/543/635/MGP01-350-700) mounted in parallel to separate different colors onto different regions of the SLM and recombine them afterwards. A transform lens (FL=400-mm) routes the light reflecting from the SLM to the photomask.

### Fluorescent collagen gels preparation

We adapted the protocol developed by Doyle et al.^37^. First, we diluted 10-mg/mL of rat tail collagen (Advanced Biomatrix, IKD119261001) with 10× reconstitution buffer, made with sodium bicarbonate (Sigma Aldrich, S5761-500G), HEPES (Sigma Aldrich, H4034-100G) and DMEM (Sigma Aldrich, D2429-100ML). This resulted in a collagen with a concentration of 3mg/mL. We then titrated the collagen to obtain a 7.4 pH. Next, we transferred ∼150-µL of the diluted pH=7.4 collagen onto 25-mm diameter #1.5 thickness coverslips (Thorlabs, CG15XH) for polymerization at room temperature. Afterwards, we incubated for 20 minutes the polymerized collagen gels with Alexa Fluor 488 NHS Ester (Thermo Fisher, A20000), which is prepared with 50-mM borate buffer (Sigma Aldrich, B6768-500G). Last, we carefully washed the sample with PBS++ (Thermo Fisher, 14040117) 3 times. Imaging experiment happened immediately after the washing step.

### Photobleaching experiment and analyses

We parked the light sheets on a single plane, tuned the intensities of both PEARLS and LLS to have similar starting maximum pixel intensities (∼17,000 camera counts or ∼9,500 photons) and kept imaging the plane with an exposure time of 10-ms for 20,000 frames. We normalized each image stack by the maximum pixel intensities of the starting frame to obtain the photobleaching curve of the stack. We then calculated the mean photobleaching curve and the standard deviation from the five locations. Last, we fit the mean photobleaching curve with an exponential decay to obtain the characteristic decay constant.

### Cell culture and imaging

HeLa cells stably expressing mCherry-CAAX and LLC-PK1 cells stably expressing mEmerald-ER and mCherry-H2B are gift from the Betzig lab at Howard Hughes Medical Institute Janelia Research Campus. All cells were cultured on pre-cleaned 25 mm diameter #1.5 coverslips coated with 10 mg/ml of fibronectin (Millipore, FC010) to aid cell adhesion. Cells were stabilized and cultured on the coverslips for 24 hours before the imaging sessions. Cells were tested for mycoplasma contamination. We performed live imaging at 37 °C, 5% CO_2_ in DMEM supplemented with 10% fetal bovine serum (Seradigm).

### Zebrafish

Zebrafish (*Danio Rerio*) were raised, and experiments were performed following regulations and protocols approved by the Institutional Animal Care and Use Committee (IACUC) at Princeton University. The zebrafish line used in this experiment is *Tg(h2az2a:h2az2a-GFP)kca66Tg* with cell nucleus ubiquitously labeled by GFP through every developmental stage. Zebrafish embryos were dechorionated manually using sharp forceps, and were mounted for imaging at the developmental stages ∼3 hours post fertilization (hpf) in 0.5% low-melting agarose (Sigma Aldrich, A9914-50G) on 25 mm diameter #1.5 thickness coverslips (Thorlabs, CG15XH). During imaging, embryos were kept in E3 embryo medium at 22°C room temperature. Tricaine methanesulfonate (MS-222) was added to the E3 medium at a concentration of 100 mg/L to immobilize the embryos at later developmental stages.

#### Drosophila melanogaster

*Drosophila* embryos co-expressing histone RFP and nascent *hunchback* (*hb*) transcripts were used from previous work^47^. Female virgins of a fly line with the fluorophores (*yw; His2Av-mRFP; nanos>NLS-MCP-mNeonGreen*) were crossed with wild-type (Ore-R) flies to reduce the background fluorescence. Female offspring (*yw/+; His2Av-mRFP/+; nanos>MCP-NLS-MCP-mNeonGreen/+*) were then crossed with males expressing the Hunchback MS2 reporter construct (48× MS2-*lacZ*), as described in ^47^. The resulting embryos from this cross were collected and used for imaging. A 25-mm in diameter #1.5 thickness coverslips (Thorlabs, CG15XH) was coated with heptane glue, made by dissolving double-sided scotch tape in heptane. The coverslip was left to dry for ∼10-min while embryos were collected, leaving an adhesive surface. Flies were caged on agar plates for 2 to 2.5-hour, and the laid embryos were transferred to a piece of double-sided tape with a dissection needle, hand-dechorionated, and placed on the glued glass coverslip. After mounting, we immersed embryos in PBS, and live imaging began at the onset of gastrulation during nuclear cycle 14.

### Image processing and visualization

Image processing were done using the GPU-based CUDA code developed by the Betzig group^5^. For fluorescent collagen gel and biological images, we applied Richard-Lucy deconvolution with the corresponding experimentally measured overall PSFs for each light sheet. 15 iterations were used for both PEARLS and LLS datasets. For image stacks that were taken using the sample stage scan mode, the raw experimental images are first deskewed before deconvolution. All visualization and rendering were prepared with napari^61^.

### Data availability

All data in the manuscript are available upon reasonable request.

## Acknowledgements

We thank T. Gregor and S. Ryabichko for the *Drosophila* line, J. Fleischer and J. Thompson for helpful discussions, the Laboratory Animal Resources at Princeton University for their help in zebrafish maintenance. T.-M.F. acknowledges the supports from National Institutes of Health (R21EB035681) and grants from the School of Engineering and Applied Science and Princeton Catalysis Initiative, Princeton University. S.K. acknowledges the support from Gilbert Omenn, M.D., Ph.D. ’61 & Martha Darling *70 Fellowship. V.G. acknowledges the support from American Heart Association Predoctoral Fellowship (24PRE1198653). C.B.Y. acknowledges the support from National Institutes of Health (NIGMS-T32GM148739). R.D.B. acknowledges the support from New Jersey Commission on Cancer Research (COCR24PRG007). K.K.-D. acknowledges the supports from Robert H. Dicke Fellowship in Experiment Physics, the National Science Foundation through the Center for the Physics of Biological Function (PHY-1734030), and Princeton University’s Dean’s for Research Innovation Fund for New Ideas.

## Author Contributions

T.-M.F. conceived the idea. Y.Q., J.Z., M.J.L., and T.-M.F. performed the theory and simulation. Y.Q., J.Z., C.R.W., and T.-M.F. built and characterized the microscope. Y.Q. and J.Z. prepared the fluorescent beads, collagen fiber, and cells. S.K., V.Z., C.B.Y., F.L., and R.D.B. prepared the zebrafish. K.K.-D. prepared the Drosophila. Y.Q., J.Z., and T.-M.F. acquired and analyzed the data, prepared the figures and wrote the manuscript with input from all authors. T.-M.F. supervised the study.

## Additional Information

Correspondence and requests for materials should be address to Tian-Ming Fu

The Supplementary Information includes (1) Supplementary Notes 1-4; (2) Table S1; (3) Figure S1-12; (4) Legends for Videos S1-8; (5) Supplementary References.

