## Supplementary Materials for "Polarization-engineered aberration-resilient light sheet microscopy"


**The PDF file includes:**

Supplementary Notes 1-3

Table S1

Figure S1-12

Legends for Videos S1-8

Supplementary References

**Other Supplementary Materials for this manuscript include the following:**

Videos S1-8

### Supplementary Notes

#### Note 1: Monochromatic nondiffracting beams

Diffraction is a fundamental feature of light beams propagating in free space, and places fundamental limits on various optical applications, including microscopy, photography, lithography, and optical communications. There has been a long-standing goal to achieve "nondiffracting" beams whose transverse spatial profiles change minimally during propagation[1]. Multiple classes of monochromatic nondiffracting beams with two-dimensional (2D) transverse spatial profiles have been theoretically predicted and experimentally realized, including Bessel, Mathieu, Weber beams and others[2]. In contrast, there are only two classes of monochromatic nondiffracting beams with 1D transverse cross-sectional profiles (i.e. light sheet): the cosine standing wave (CSW) and the Airy beam[1]. The 1D Airy beam is generally unsuitable for light sheet applications because its center of mass shifts parabolically during propagation[3] (Figure 1G). In contrast, CSW maintains a fixed beam center while preserving a propagation invariant transverse profile (Figure S1F).

To clarify the terminology used in this work, we define the following categories of light sheet profiles. A light sheet is spatiotemporally invariant if its profile remains unchanged along the propagation direction (i.e., nondiffracting) and remains static in time without requiring either spatial or temporal averaging. It is spatially invariant but temporally variant if its profile remains unchanged along the propagation direction but requires scanning, as in Bessel beams, lattice light sheet (LLS), or Field Synthesis light sheet. It is temporally invariant but spatially variant if its profile changes along the propagation direction but does not require scanning, as in a Gaussian light sheet generated by a cylindrical lens. Finally, a beam is spatiotemporally variant if its profile changes during propagation and scanning is required, as in a scanning Gaussian light sheet.

#### Note 2: Nondiffracting condition of CSWs

For CSW, we define the propagation length (i.e., “nondiffracting” region), as the full width at half maximum (FWHM) of the beam intensity along the y-axis. Analytically, it can be expressed as[4]:

$$y_{FWHM} \approx \frac{\pi}{[(\boldsymbol{k}_{min} - \boldsymbol{k}_{max}) \cdot \hat{\boldsymbol{e}}_y]} = \frac{\lambda_{illumination}/n}{2(\sqrt{1 - \left(\frac{NA_{min}}{n}\right)^2} - \sqrt{1 - \left(\frac{NA_{max}}{n}\right)^2})}$$

In order for all the constituent CSWs of PEARLS to have the same propagation length, the following nondiffracting condition must be satisfied:

$$\sqrt{1 - \left(\frac{NA_k^{min}}{n}\right)^2} - \sqrt{1 - \left(\frac{NA_k^{max}}{n}\right)^2} = constant$$

where $NA_k^{max}$ and $NA_k^{min}$ are the maximum and minimum numerical apertures (NAs) of the $k^{\text{th}}$ CSW, respectively. Under paraxial approximation, $\sqrt{1 - \left(\frac{NA_k^{min}}{n}\right)^2} \approx 1 - \frac{1}{2}\left(\frac{NA_k^{min}}{n}\right)^2$ and $\sqrt{1 - \left(\frac{NA_k^{max}}{n}\right)^2} \approx 1 - \frac{1}{2}\left(\frac{NA_k^{max}}{n}\right)^2$. Therefore, the nondiffracting condition can be expressed as:

$${NA_k^{max}}^2 - {NA_k^{min}}^2 = (NA_k^{max} + NA_k^{min})(NA_k^{max} - NA_k^{min}) = constant$$

**Note 3: PEARLS can produce any types of dithered LLS**

Based on crystallography, there are only five 2D Bravais lattices: oblique, rectangular, centered rectangular, square, and hexagonal. Among these, only hexagonal and square lattices were selected for LLS microscopy thanks to their high axial resolution and transverse symmetry[5]. The corresponding optical transfer functions (OTFs) of ideal dithered hexagonal and square lattices with infinite long propagation are[4,5]:

- Hexagonal LLS:

$$OTF_{LLS}^{HEX}(k_z) = \delta(k_z) + \frac{\delta(k_z \pm k_0 NA)}{3} + \frac{\delta(k_z \pm 2k_0 NA)}{6}$$

- Square LLS:

$$OTF_{LLS}^{SQ}(k_z) = \delta(k_z) + \frac{\delta(k_z \pm 2k_0 NA)}{4}$$

On the other hand, the OTF of an ideal CSW with infinite long propagation is[4]:

$$OTF_{CSW}(k_z) = \delta(k_z) + \frac{\delta(k_z \pm 2k_0 NA)}{2}$$

Since PEARLS is the incoherent summation of two CSWs, its OTF can be written as:

$$OTF_{PEARLS}(k_z) = (\alpha + \beta)\delta(k_z) + \alpha\frac{\delta(k_z \pm 2k_0 NA^p)}{2} + \beta\frac{\delta(k_z \pm 2k_0 NA^s)}{2}$$

where $NA^p$ and $NA^s$ is the NAs of the p- and s-polarized CSW, respectively, and $\alpha$ and $\beta$ is the relative intensity between the p- and s-polarized CSW, respectively.

- To produce the equivalent spatial profile of dithered hexagonal LLS, we set $NA^s = 2NA^p = NA$, $\alpha = 2/3$, and $\beta = 1/3$, which leads:

$$OTF_{PEARLS}^{HEX}(k_z) = \delta(k_z) + \frac{\delta(k_z \pm k_0 NA)}{3} + \frac{\delta(k_z \pm 2k_0 NA)}{6} = OTF_{LLS}^{HEX}(k_z)$$

- To produce the equivalent spatial profile of dithered square LLS, we set $NA^p = 0$ and $NA^s = NA$, and $\alpha = \beta = 1/2$, which gives:

$$OTF_{PEARLS}^{SQ}(k_z) = \delta(k_z) + \frac{\delta(k_z \pm 2k_0 NA)}{4} = OTF_{LLS}^{HEX}(k_z)$$

### Supplementary Table

### Table S1 | Imaging conditions

| Figures and Videos | Specimen | Illumination mode | Exposure (ms) | Laser (nm) | Scan mode | Scan range (μm) | Scan step (nm) | Time interval (s) |
|---|---|---|---|---|---|---|---|---|
| Figure 2E, Video S3 | Collagen gel | LLS (Hexagonal, NA 0.6/0.5)<br>PEARLS (Hexagonal, NA 0.6/0.5) | 10 | 488 | N/A | N/A | N/A | 0.01 |
| Figure 2I | Collagen gel | LLS (Square, NA 0.6/0.5)<br>PEARLS (Square, NA 0.6/0.5) | 100 | 488 | Objective scan | 10 | 100 | N/A |
| Figure S9D,E | Collagen gel | PEARLS (Square, NA 0.6/0.5) | 100 | 488 | Sample scan | 100 | 200 | N/A |
| Figure S9F,G | Collagen gel | LLS (Square, NA 0.6/0.5)<br>PEARLS (Square, NA 0.6/0.5) | 100 | 488 | Objective scan | 20 | 100 | N/A |
| Figure S9I | Collagen gel | LLS (Square, NA 0.6/0.5)<br>PEARLS (Square, NA 0.6/0.5) | 100 | 488 | N/A | N/A | N/A | 0.1 |
| Video S4 | Collagen gel | LLS (Square, NA 0.6/0.5)<br>PEARLS (Square, NA 0.6/0.5) | 10 | 488 | N/A | N/A | N/A | 0.01 |
| Figure 3A-C, Video S5 | HeLa | PEARLS (Square, NA 0.5/0.4) | 50 | 560 | Sample scan | 100 | 400 | 13.2 |
| Figure 3D-F, Video S6 | LLC-PK1 | PEARLS (Square, NA 0.5/0.4) | 50 | 488 & 560 | Sample scan | 110 | 400 | 15.7 |
| Figure 4A,B, Video S7 | Zebrafish | PEARLS (Square, NA 0.42/0.38) | 40 | 488 | Sample scan | 250 | 400 | 26.8 |
| Figure S11A | Zebrafish | PEARLS (Square, NA 0.55/0.50) | 100 | 488 | Sample scan | 100 | 400 | 25.1 |
| Figure S11B | Zebrafish | LLS (Square, NA 0.55/0.50)<br>PEARLS (Square, NA 0.55/0.50) | 100 | 488 | Sample scan | 100 | 400 | 25.1 |

| | | | | | | | | |
|---|---|---|---|---|---|---|---|---|
| Figure 4C,D, Video S8 | Drosophila | PEARLS (Square, NA 0.42/0.38) | 100 | 488 & 560 | Sample scan | 450 | 400 | 123 |

### Supplementary Figures

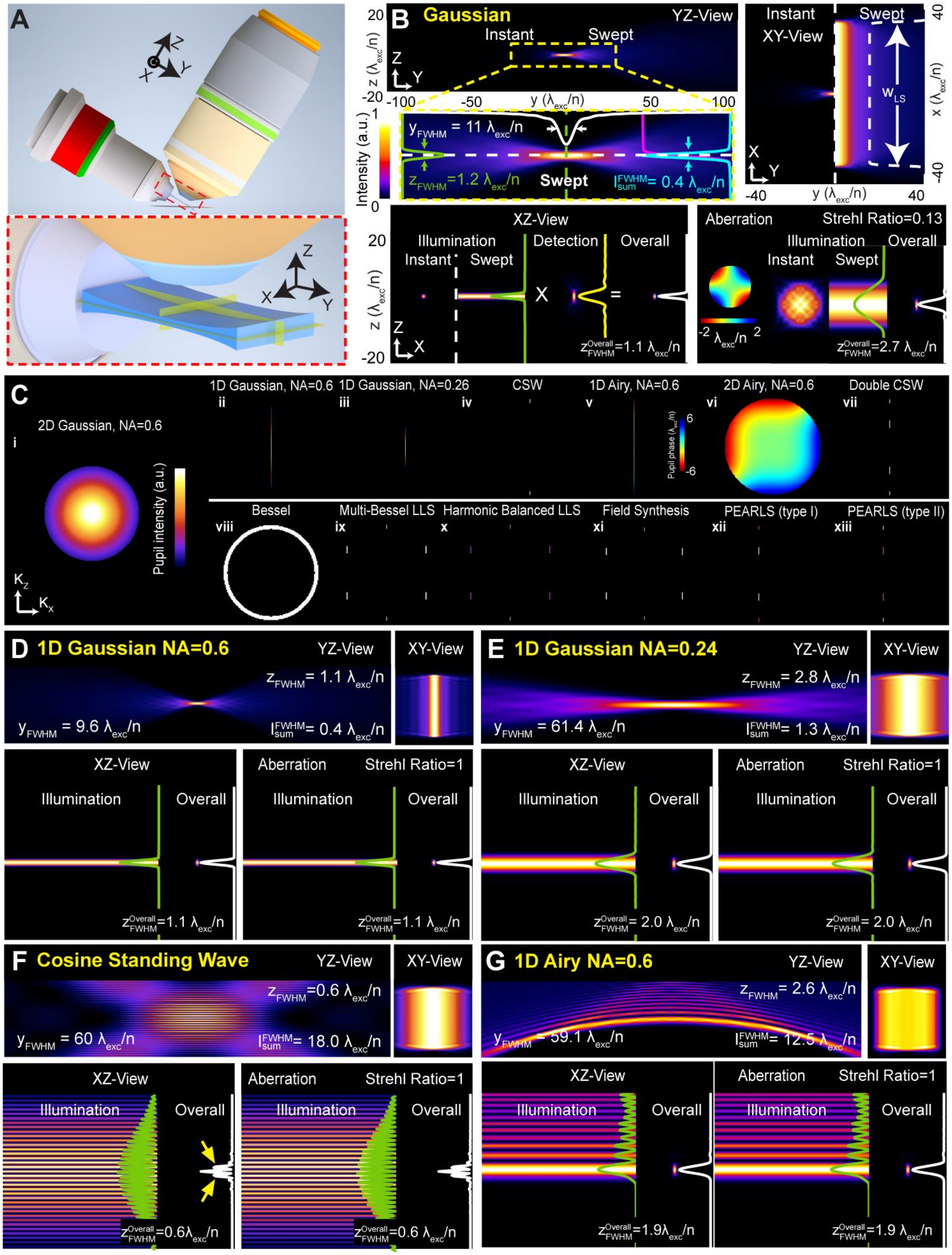

**Figure S1 | Comparisons of different types of light sheets (LSs).** (A) Objective configuration and coordinate definition. X, Y, and Z axes indicate the LS width, propagation, and thickness directions, respectively. (B) Example characterizations of a 2D scanning Gaussian LS (NA=0.6). Multiple metrics are presented, including instant and swept/scanned (if applicable) YZ and XY profiles of the LS. In addition, instant and swept/scanned (if applicable) XZ profiles with their Z-axis line cuts, as well as the overall XZ PSFs with corresponding Z-axis line cuts, are shown for both cases with and without astigmatism aberration. Quantified parameters include the propagation length ($Y_{\mathrm{FWHM}}$), axial confinement ($I_{Sum}^{FWHM}$, defined as the axial FWHM of the accumulated energy), axial resolution of the LS ($Z_{\mathrm{FWHM}}$), axial resolution of the overall PSF ($Z_{FWHM}^{Overall}$), and Strehl ratio under astigmatism aberration. Desirable performance corresponds to larger $Y_{\mathrm{FWHM}}$, lower $I_{Sum}^{FWHM}$, lower $Z_{\mathrm{FWHM}}$, lower $Z_{FWHM}^{Overall}$, and higher Strehl ratio. (C) Electric field intensity on the pupils of various LSs. (D-G) Characterizations of non-scanning 1D Gaussian LS (D, NA=0.6), non-scanning 1D Gaussian LS (E, NA=0.24), cosine standing wave (CSW, F), 1D Airy LS (G, NA=0.6). For each case, YZ and XY profiles of the LS are shown, as well as the XZ profiles of the LS and overall PSFs, evaluated with and without applying the astigmatism aberration shown in (B).

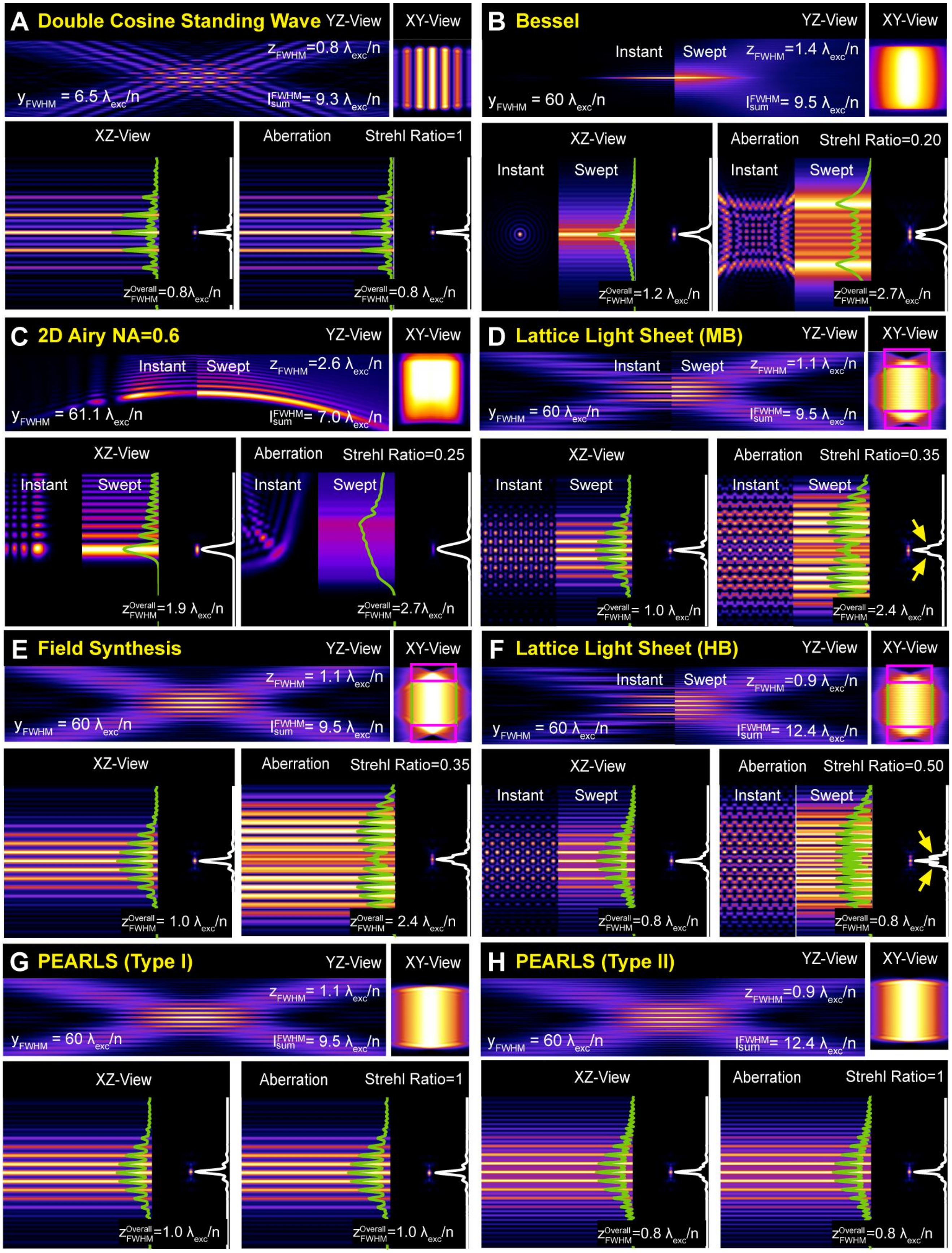
A Double Cosine Standing Wave
YZ-View
XY-View
$z_{FWHM}$=0.8 $\lambda_{exc}$/n
$y_{FWHM}$ = 6.5 $\lambda_{exc}$/n
$I_{sum}^{FWHM}$= 9.3 $\lambda_{exc}$/n
XZ-View
Aberration
Strehl Ratio=1
$z_{FWHM}^{Overall}$=0.8$\lambda_{exc}$/n
$z_{FWHM}^{Overall}$=0.8 $\lambda_{exc}$/n
B Bessel
YZ-View
XY-View
Instant
Swept
$z_{FWHM}$=1.4 $\lambda_{exc}$/n
$y_{FWHM}$ = 60 $\lambda_{exc}$/n
$I_{sum}^{FWHM}$= 9.5 $\lambda_{exc}$/n
XZ-View
Aberration
Strehl Ratio=0.20
Instant
Swept
$z_{FWHM}^{Overall}$=1.2 $\lambda_{exc}$/n
Instant
Swept
$z_{FWHM}^{Overall}$=2.7$\lambda_{exc}$/n
C 2D Airy NA=0.6
YZ-View
XY-View
Instant
Swept
$z_{FWHM}$=2.6 $\lambda_{exc}$/n
$y_{FWHM}$ = 61.1 $\lambda_{exc}$/n
$I_{sum}^{FWHM}$= 7.0 $\lambda_{exc}$/n
XZ-View
Aberration
Strehl Ratio=0.25
Instant
Swept
$z_{FWHM}^{Overall}$=1.9 $\lambda_{exc}$/n
Instant
Swept
$z_{FWHM}^{Overall}$=2.7$\lambda_{exc}$/n
D Lattice Light Sheet (MB)
YZ-View
XY-View
Instant
Swept
$z_{FWHM}$=1.1 $\lambda_{exc}$/n
$y_{FWHM}$ = 60 $\lambda_{exc}$/n
$I_{sum}^{FWHM}$= 9.5 $\lambda_{exc}$/n
XZ-View
Aberration
Strehl Ratio=0.35
Instant
Swept
$z_{FWHM}^{Overall}$= 1.0 $\lambda_{exc}$/n
Instant
Swept
$z_{FWHM}^{Overall}$=2.4 $\lambda_{exc}$/n
E Field Synthesis
YZ-View
XY-View
$z_{FWHM}$= 1.1 $\lambda_{exc}$/n
$y_{FWHM}$ = 60 $\lambda_{exc}$/n
$I_{sum}^{FWHM}$= 9.5 $\lambda_{exc}$/n
XZ-View
Aberration
Strehl Ratio=0.35
$z_{FWHM}^{Overall}$= 1.0 $\lambda_{exc}$/n
$z_{FWHM}^{Overall}$= 2.4 $\lambda_{exc}$/n
F Lattice Light Sheet (HB)
YZ-View
XY-View
Instant
Swept
$z_{FWHM}$=0.9 $\lambda_{exc}$/n
$y_{FWHM}$ = 60 $\lambda_{exc}$/n
$I_{sum}^{FWHM}$= 12.4 $\lambda_{exc}$/n
XZ-View
Aberration
Strehl Ratio=0.50
Instant
Swept
$z_{FWHM}^{Overall}$=0.8 $\lambda_{exc}$/n
Instant
Swept
$z_{FWHM}^{Overall}$=0.8 $\lambda_{exc}$/n
G PEARLS (Type I)
YZ-View
XY-View
$z_{FWHM}$= 1.1 $\lambda_{exc}$/n
$y_{FWHM}$ = 60 $\lambda_{exc}$/n
$I_{sum}^{FWHM}$= 9.5 $\lambda_{exc}$/n
XZ-View
Aberration
Strehl Ratio=1
$z_{FWHM}^{Overall}$=1.0 $\lambda_{exc}$/n
$z_{FWHM}^{Overall}$=1.0 $\lambda_{exc}$/n
H PEARLS (Type II)
YZ-View
XY-View
$z_{FWHM}$=0.9 $\lambda_{exc}$/n
$y_{FWHM}$ = 60 $\lambda_{exc}$/n
$I_{sum}^{FWHM}$= 12.4 $\lambda_{exc}$/n
XZ-View
Aberration
Strehl Ratio=1
$z_{FWHM}^{Overall}$=0.8 $\lambda_{exc}$/n
$z_{FWHM}^{Overall}$=0.8 $\lambda_{exc}$/n

**Figure S2 | Comparisons of different types of LSs (continued).** (A) double CSW LS with two coherently superposed CSWs. (B) Scanning Bessel LS. (C) Scanning 2D Airy beam. (D) Multi-Bessel lattice light sheet (MB-LLS). (E) Field synthesis LS. (F) Harmonic balanced lattice light sheet (HB-LLS). (G) Type I PEARLS with similar profiles of MB-LLS. (H) Type II PEARLS with similar profiles of HB-LLS.

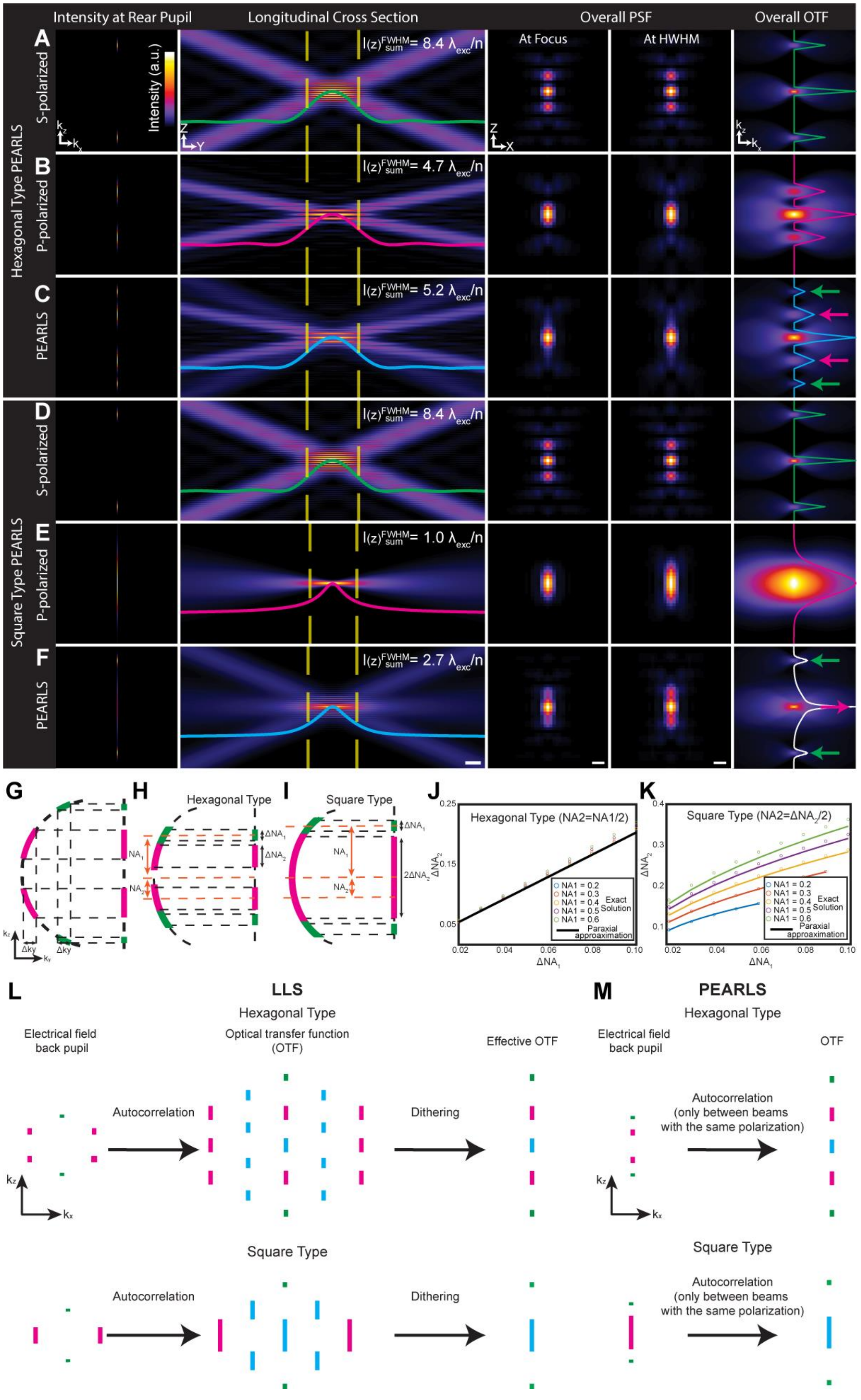

Intensity at Rear Pupil
Longitudinal Cross Section
Overall PSF
Overall OTF
At Focus
At HWHM
Hexagonal Type PEARLS
Square Type PEARLS
A
S-polarized
B
P-polarized
C
PEARLS
D
S-polarized
E
P-polarized
F
PEARLS
Intensity (a.u.)
I(z)FWHM sum= 8.4 λexc/n
I(z)FWHM sum= 4.7 λexc/n
I(z)FWHM sum= 5.2 λexc/n
I(z)FWHM sum= 8.4 λexc/n
I(z)FWHM sum= 1.0 λexc/n
I(z)FWHM sum= 2.7 λexc/n
G
H
Hexagonal Type
I
Square Type
J
Hexagonal Type (NA2=NA1/2)
K
Square Type (NA2=ΔNA2/2)
NA1 = 0.2
NA1 = 0.3
NA1 = 0.4
NA1 = 0.5
NA1 = 0.6
Exact Solution
Paraxial approaximation
L
LLS
Hexagonal Type
Electrical field back pupil
Optical transfer function (OTF)
Effective OTF
Autocorrelation
Dithering
Square Type
Autocorrelation
Dithering
M
PEARLS
Hexagonal Type
Electrical field back pupil
OTF
Autocorrelation (only between beams with the same polarization)
Square Type
Autocorrelation (only between beams with the same polarization)

**Figure S3 | Nondiffracting condition and universal generation of dithered LLS.** (A-C) A hexagonal type PEARLS (C) consists of two CSWs—one with s-polarization (A) and one with p-polarization (B)—that have the same propagation length. (D-F) A square type PEARLS (F) consists of two orthogonally polarized CSWs that have the same propagation length. (A-F) For each row, columns from left to right: intensity distribution at the rear pupil. YZ cross-sectional profiles. Curves showing the corresponding intensity profiles on the Y-axis. Yellow dashed lines indicate HWHMs of the intensity profiles and define the nondiffracting region (Scale bar: 10-$\lambda_{exc}$/n). The corresponding axial confinement values $I(z)_{Sum}^{FWHM}$ are labeled at the upper right of the YZ cross-sectional profiles. XZ cross-sectional profiles of the overall PSFs at the foci (third column) and at HWHM (fourth column) of the light sheets. Scale bars: 1-$\lambda_{exc}$/n. Fifth column: XZ cross-sectional profiles of the overall OTFs with line cuts along $k_z$. Colored arrows indicate spatial frequency contributions from the individual CSW component. (G) Schematic illustration of the nondiffracting condition on the rear pupil space: two CSWs (magenta and green) need to have the same momentum spreading along the propagation direction on the Ewald sphere to have identical propagation length. Projections of these wavevectors from the Ewald sphere to the rear pupil plane set the nondiffracting condition. (H and I) Schematics showing nondiffracting condition in (G) applied to hexagonal-type (H) and square-type (I) PEARLS configuration. Yellow and black arrows indicate NA and ΔNA, respectively. (J and K) Comparisons between the calculated ΔNA values from the exact nondiffracting condition and the paraxial approximation. Simulations were performed with $NA_1$ ranging from 0.2 to 0.6 and $\Delta NA_1$ from 0.02 to 0.10. Circles represent the exact solution, and lines show the paraxial approximation. (L) Electrical fields at the back pupil (left), OTFs (middle), effective OTFs after dithering (right) of hexagonal (top) and square (bottom) LLS. Green and magenta lines represent correlation of beams with the same $|k_x|$, and blue lines showing cross terms between beams with different

$|k_x|$. (M) Electrical fields and OTFs of corresponding PEARLS. Autocorrelation happens only between beams with the same polarization. Green and magenta lines indicate the two incoherent CSWs.

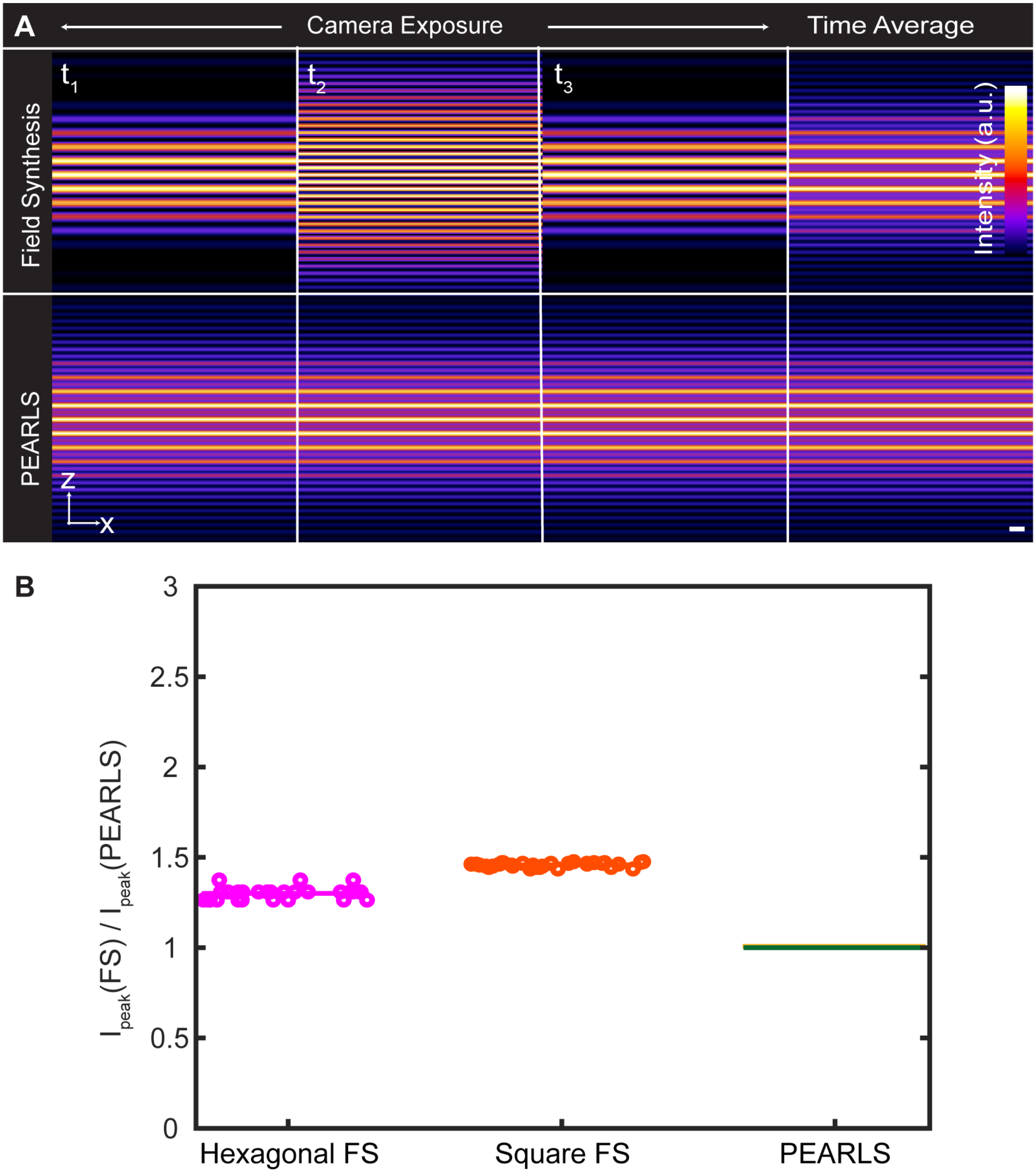


**Figure S4 | Comparisons between Field Synthesis (FS) and PEARLS.** (A) Instantaneous (column 1-3) and time averaged (column 4) XZ transverse profiles of FS (top) and PEARLS (bottom) within a single exposure. Unlike FS that has a temporally varying profile, PEARLS maintains a constant illumination profile. Scale bar: 2.5-$\lambda_{exc}$/n. (B) Simulated peak illumination power ratios between FS and corresponding PEARLS with the same transverse cross-section profiles and resulted image SNR. Each group contains various combinations of NA and ΔNA, ranging from 0.3-0.6 and 0.02-0.2, respectively.

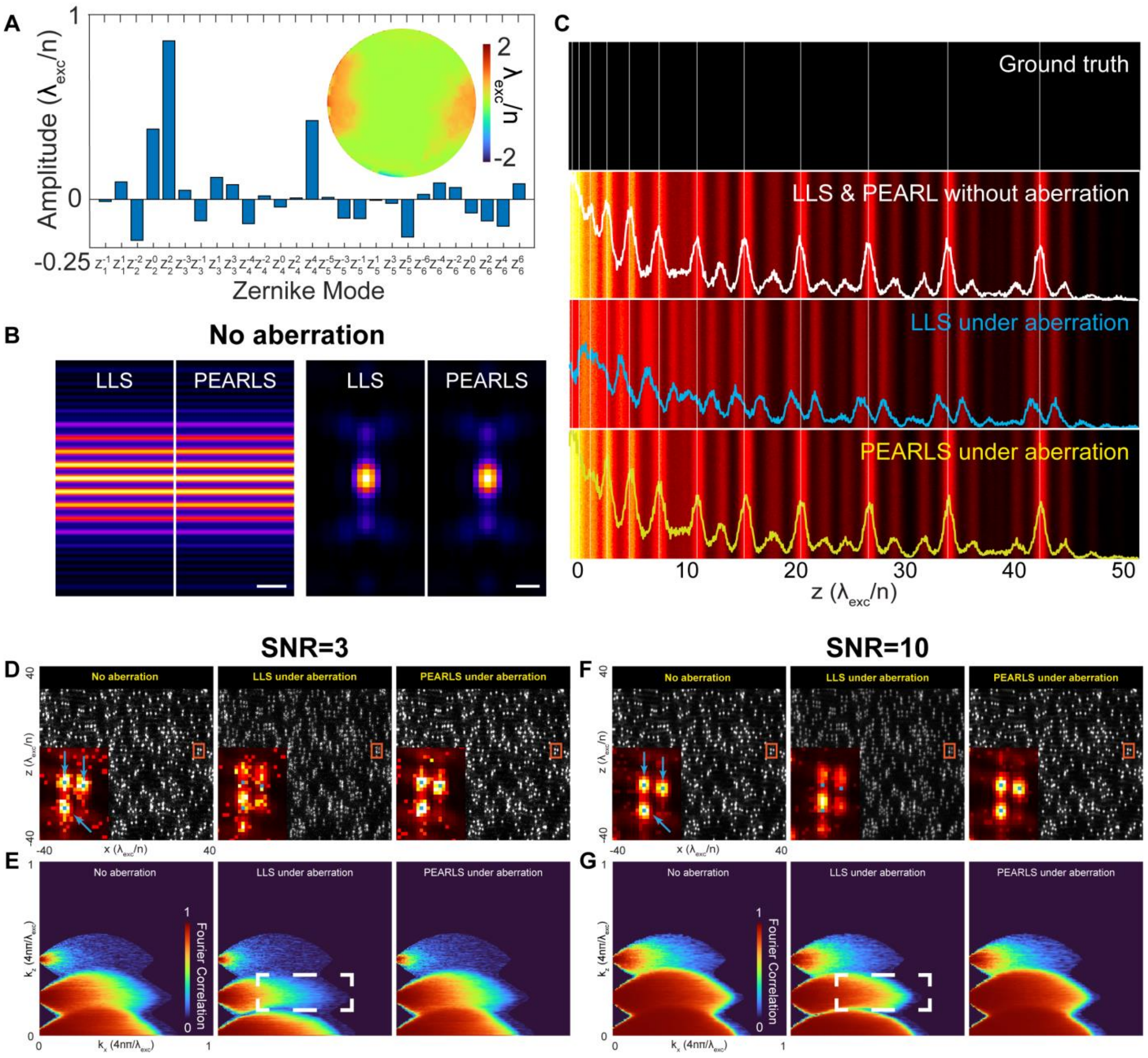


**Figure S5 | Characterizations of PEARLS and LLS under measured aberration from zebrafish spinal cord.** (A) Decomposition of the aberration into Zernike modes. (B) Transverse cross-section profiles and overall PSFs of LLS and PEARLS without aberration. Scale bars: 5-$\lambda_{exc}$/n (cross-section profiles) and 1-$\lambda_{exc}$/n (overall PSF). (C) Axial resolution characterizations of PEARLS and LLS without (second row) and with (third row: LLS; fourth row: PEARLS) the aberration applied using a line pattern with variable spacings. The white, blue, and yellow curves are corresponding line cuts. SNR=10 was used for all simulations. (D) Simulated images of point emitters imaged by LLS and PEARL without (left) and with (middle: LLS; right: PEARLS) the aberration applied. Insets: zoom-in view of the red box highlighting three closely spaced point emitters. The three blue dots indicated by the arrows are the ground truth positions of the emitters. Simulations were performed at SNR = 3. (E) Fourier shell correlation analyses of the results in (D). The white dashed box in the middle panel highlights the origin of aberration degradation of LLS. (F and G) Same analysis as in (D and E), performed at a higher SNR of 10.

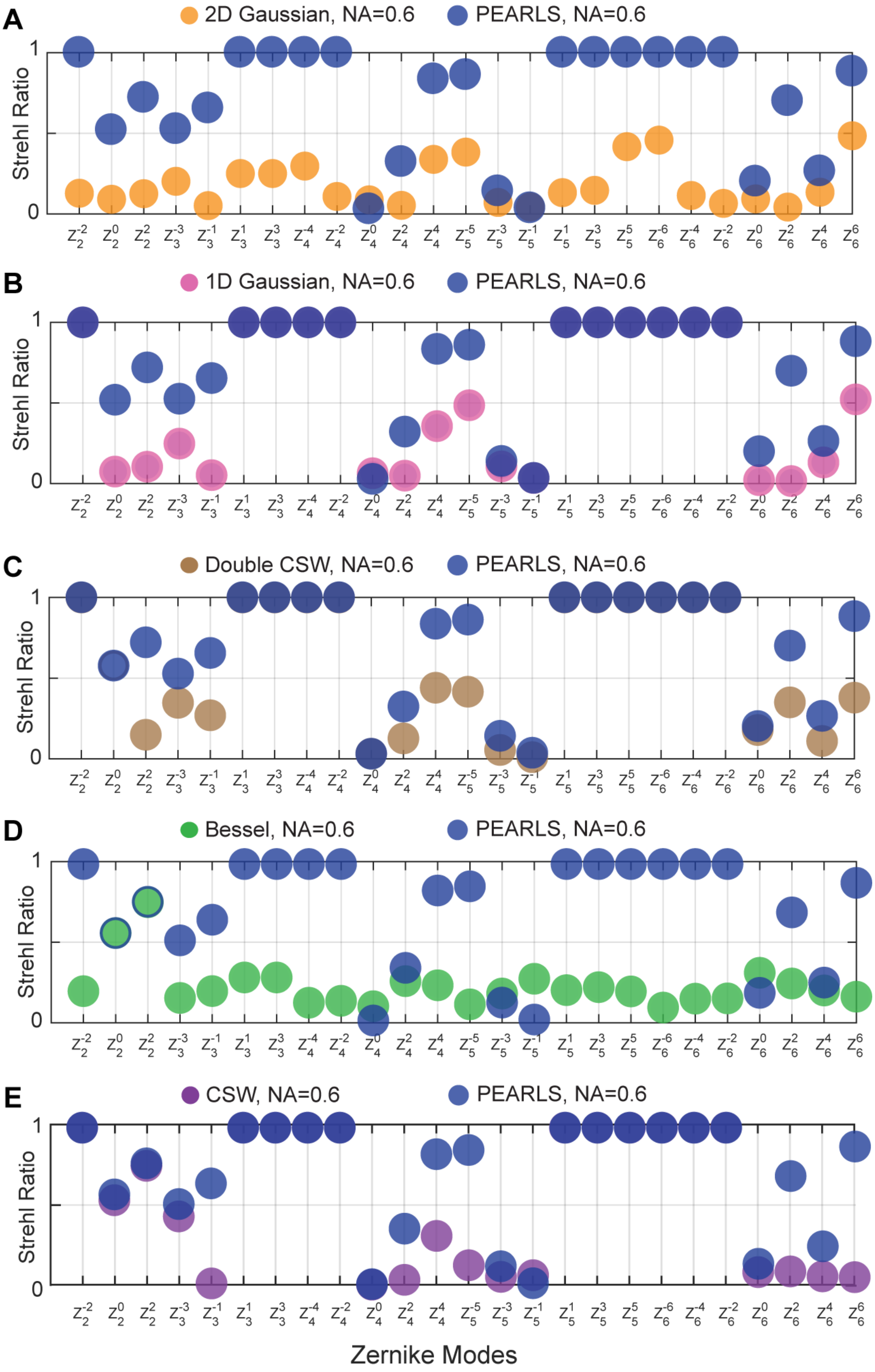
A
2D Gaussian, NA=0.6
PEARLS, NA=0.6
B
1D Gaussian, NA=0.6
PEARLS, NA=0.6
C
Double CSW, NA=0.6
PEARLS, NA=0.6
D
Bessel, NA=0.6
PEARLS, NA=0.6
E
CSW, NA=0.6
PEARLS, NA=0.6
Strehl Ratio
1
0
$Z_2^{-2}$ $Z_2^0$ $Z_2^2$ $Z_3^{-3}$ $Z_3^{-1}$ $Z_3^1$ $Z_3^3$ $Z_4^{-4}$ $Z_4^{-2}$ $Z_4^0$ $Z_4^2$ $Z_4^4$ $Z_5^{-5}$ $Z_5^{-3}$ $Z_5^{-1}$ $Z_5^1$ $Z_5^3$ $Z_5^5$ $Z_6^{-6}$ $Z_6^{-4}$ $Z_6^{-2}$ $Z_6^0$ $Z_6^2$ $Z_6^4$ $Z_6^6$
Zernike Modes

**Figure S6 | Strehl ratio comparisons between PEARLS and other light sheets.** (A-E) Strehl ratios of PEARLS (dark blue) and other LSs, including (A) scanning 2D Gaussian LS (orange), (B) non-scanning 1D Gaussian LS (magenta), (C) double CSW LS with two coherently superposed CSWs (brown), (D) scanning Bessel LS (green), and (E) single CSW LS (purple) under 25 different Zernike aberration modes**.** Gaussian LSs have a NA of 0.6, and all the other LSs have $NA_{max}/NA_{min}$=0.6/0.56.

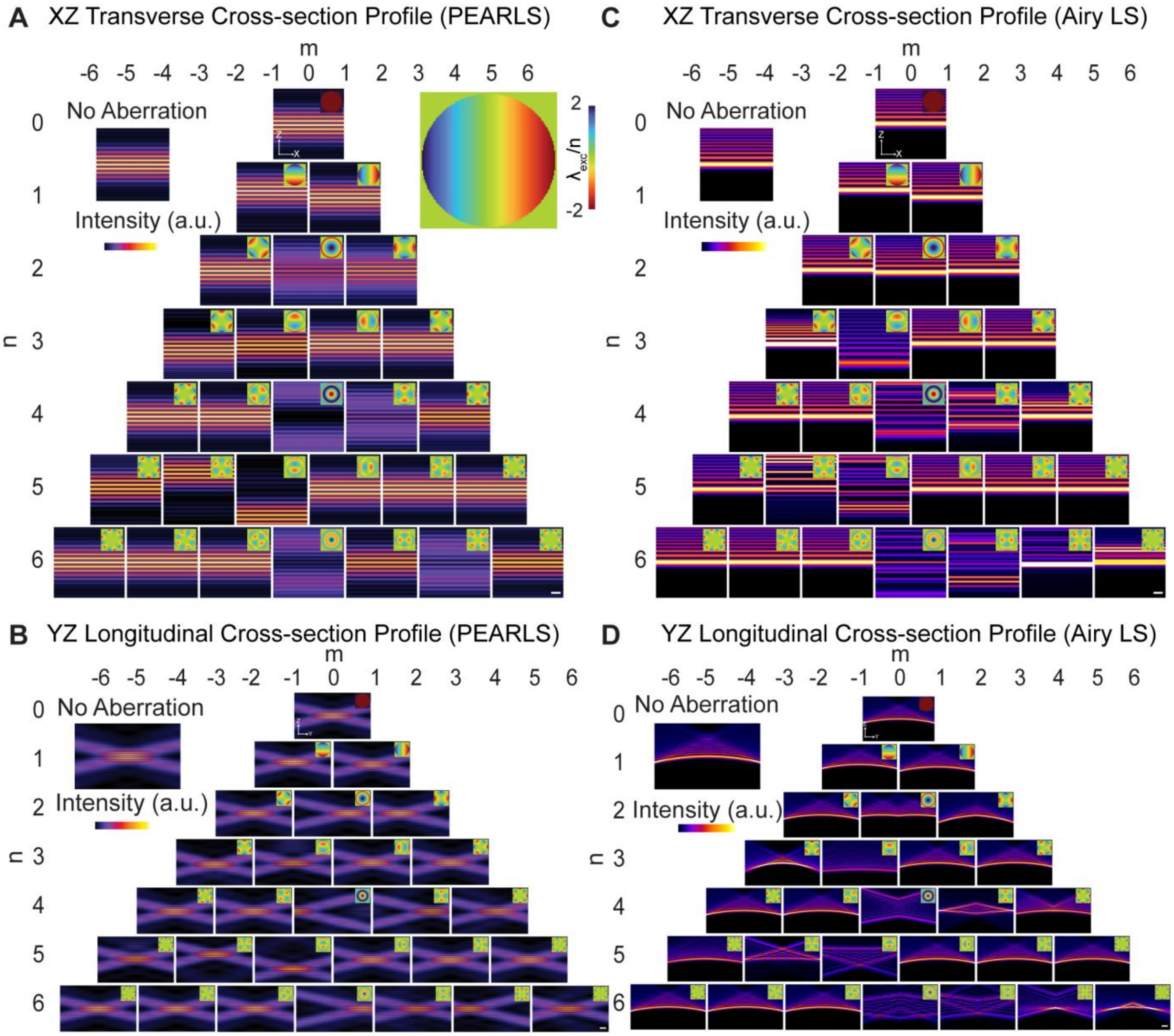


**Figure S7 | Transverse and longitudinal profiles of PEARLS and 1D Airy LS under different Zernike aberration modes.** (A and B) Transverse (XZ) and Longitudinal (YZ) profiles of PEARLS under aberrations. Each panel shows the XZ and YZ profile under a single aberration mode, with the corresponding wavefront aberration shown as an inset (colormap in units of $\lambda_{exc}/n$).

(C and D) Corresponding XZ and YZ profiles for an Airy light sheet (Airy LS) generated with the same NA as in (A) and (B) under the same aberration modes.

Scale bars: 5-$\lambda_{exc}/n$ (XZ profiles) and 10 $\lambda_{exc}/n$ (YZ profiles).

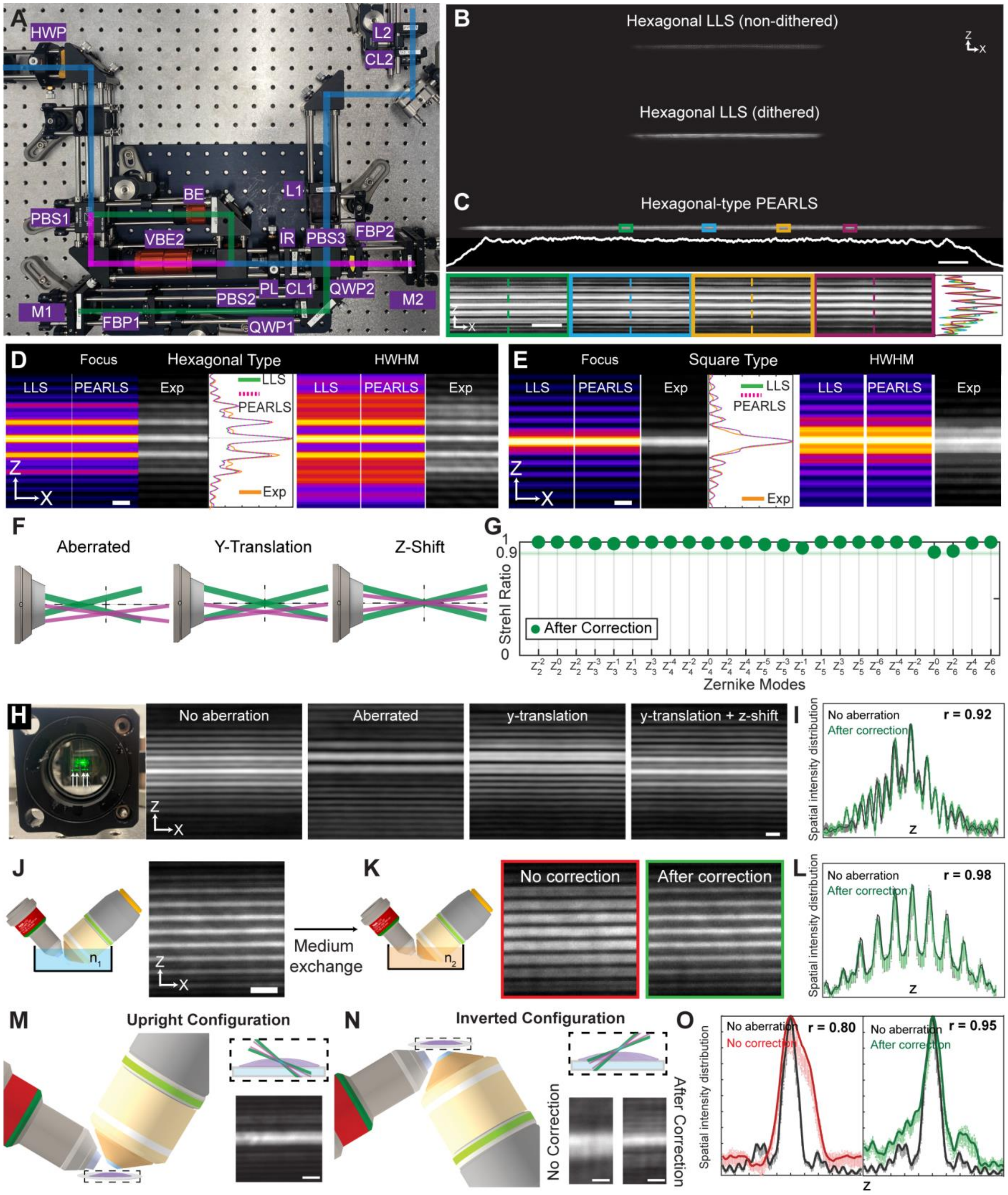
A
HWP
L2
CL2
BE
L1
PBS1
VBE2
IR
PBS3
FBP2
PL
CL1
QWP2
PBS2
M2
M1
FBP1
QWP1
B
Hexagonal LLS (non-dithered)
Hexagonal LLS (dithered)
C
Hexagonal-type PEARLS
D
Focus
Hexagonal Type
HWHM
LLS
PEARLS
Exp
E
Square Type
F
Aberrated
Y-Translation
Z-Shift
G
Strehl Ratio
After Correction
Zernike Modes
H
No aberration
Aberrated
y-translation
y-translation + z-shift
I
Spatial intensity distribution
After correction
r = 0.92
J
K
Medium exchange
No correction
After correction
L
r = 0.98
M
Upright Configuration
N
Inverted Configuration
No Correction
After Correction
O
r = 0.80
r = 0.95

**Figure S8 | Experimental realizations and characterizations of PEARLS.**

(A) Photograph of the beam shaping module. HWP, half-wave plate; PBS, polarizing beamsplitter; VBE, variable beam expander; BE, beam expander; PL, Powell lens; IR, iris; CL, cylindrical lens; L, lens; QWP, quarter-wave plate; FBP, Fresnel biprism; M, mirror. (B) XZ cross-sectional profiles of a non-dithered (top) and dithered (bottom) hexagonal LLS ($NA_{max}/NA_{min}$=0.6/0.52) with a width of ~200-µm that is limited by the width of the spatial light modulator (SLM). (C) XZ cross-sectional profile of a ~750-µm wide hexagonal-type PEARLS ($NA_{max}/NA_{min}$=0.6/0.52). The white line cut shows the uniform central intensity profile of PEARLS along the width direction (X-axis). Four color-coded boxes highlight selected regions along PEARLS, with corresponding zoomed-in views below. The plots on the far-right show the linecuts taken at the four colored dashed lines inside corresponding boxes. Scale bar: 50-µm. (D and E) XZ cross-sectional profiles and line cuts comparisons between hexagonal (D) and square (E) type LLS and PEARLS at the focal and half width HWHM planes. Scale bar: 2.5-$\lambda_{exc}$/n. (F) Schematics showing the effect of optical aberration and correction steps for PEARLS. (G) Strehl ratios of PEARLS after aberration correction steps in (F). A Strehl ratio of 0.9 (green line) is achieved under all 25 Zernike aberrations. (H) Aberration correction capability of PEARLS. Left: A plastic film mounted at a rear pupil conjugate plane introduced an optical aberration. Right: experimental results showing PEALRS transverse cross-section profiles without aberration, after aberration, after y-translation of the p-polarized CSW, and after z-shift of the p-polarized CSW, respectively. Scale bar: 2-µm. (I) Spatial intensity distributions of the aberration-free and the corrected profile with the cross-correlation coefficient (Pearson's r) indicated. (J) Aligned transverse profile of PEARLS in 1×PBS ($n_1$=1.334). Scale bar: 2-µm. (K) Aberrated (red box) and corrected profile (green box) after medium exchange to deionized water ($n_2$=1.332). (L) Spatial intensity distributions in 1×PBS and after correction in water with the cross-correlation coefficient indicated. (M) Schematic showing standard upright objectives configuration (left). Zoom-in view of PEARLS and the specimen on top of a coverslip (#1.5, ~170-µm thick) (top right). Transverse illumination profile of a square-type PEARLS (bottom right). Scale bar: 2-µm. (N) Schematic showing inverted objectives configuration (left). Zoom-in view of PEARLS that passes through the coverslip and the specimen (top right). Transverse illumination profile of the same PEARLS in (M) before (bottom right, column 1) and after (bottom right, column 2) aberration correction. Scale bar: 2-µm. (O) Spatial intensity distributions of the aberration-free, no-correction, and after-correction profiles with the cross-correlation coefficient indicated.

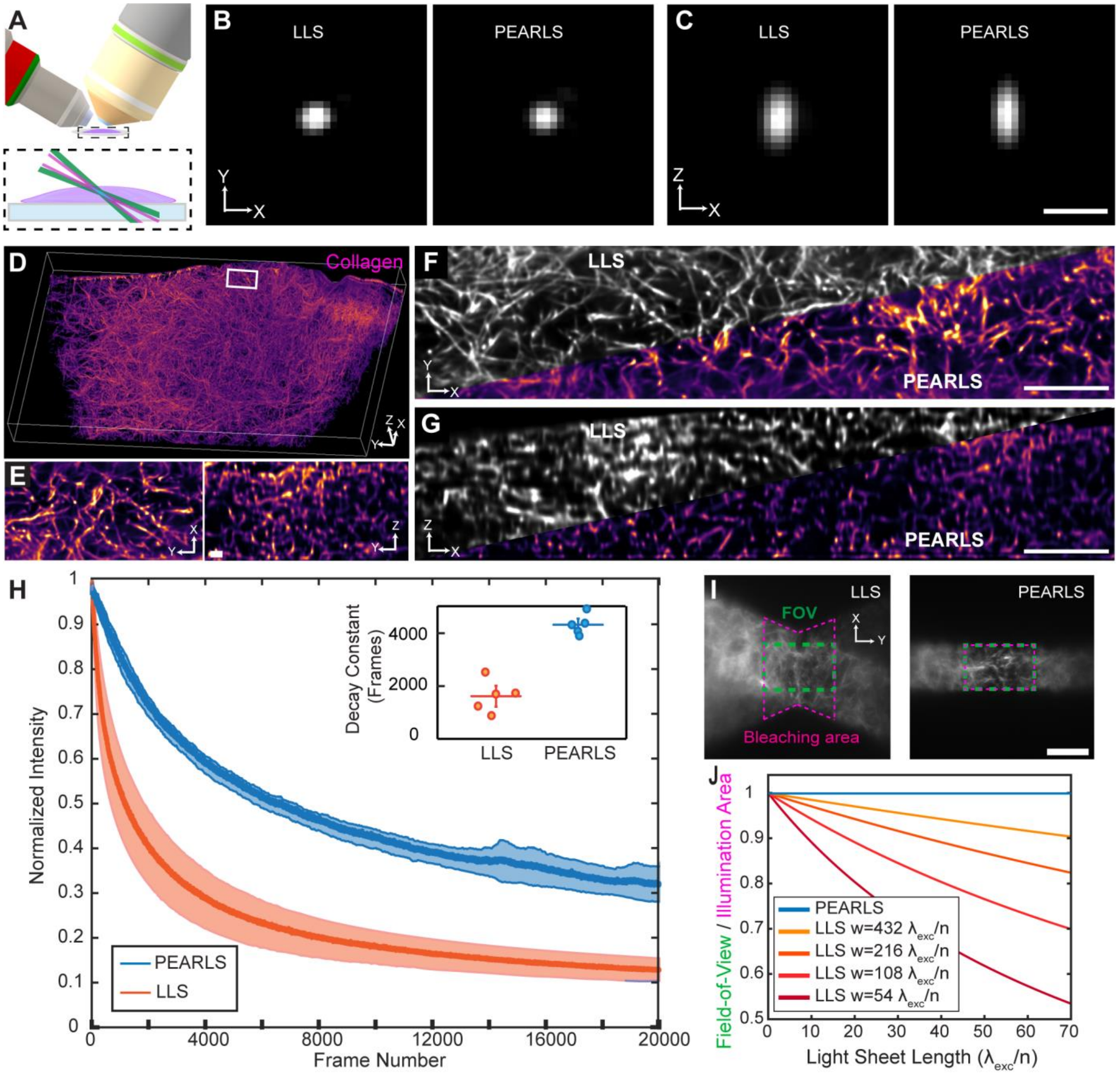


**Figure S9 | 3D imaging of fluorescent collagen gel using LLS and PEARLS.** (A) Schematic showing objective configurations (left: illumination objective; right: detection objective; bottom: specimen on a coverslip). (B and C) Lateral (B) and axial (C) slice views of the overall PSFs from LLS (left) and PEARLS (right). Scale bar: 1-µm. (D) Volume rendering of a collagen gel labeled with AlexaFluor 488 imaged using PEARLS. Bounding box: 111-µm by 194-µm by 31-µm. (E) Zoomed-in XY (left) and YZ (right) views of the white box in (D). Scale bar: 2-µm. (F and G) Comparisons of lateral (F) and axial (G) slice views of the volumetrically imaged collagen gels using LLS (gray color, top left) and PEARLS (purple color, bottom right). Scale bar: 12-µm. (H) Averaged photobleaching curves of PEARLS (blue) and LLS (red). The shaded areas indicate ±1 standard deviation (SD) from five datasets taken at five different regions of the same collagen gel. Intensities were normalized with respect to the maximum value of the corresponding Frame 1. Inset: characteristic decay constants of LLS (red) and PEARLS (blue) obtained by fitting individual photobleaching curves with exponential decays. Horizontal lines indicate average of the five datasets of each light sheet, and vertical lines indicate ±1 SD. (I) Out-of-FOV photobleaching comparison of LLS (left) and PEARLS (right) with the same cross-section profile. Raw collagen images are shown. Magenta dashed lines: bleaching regions. Green dashed line: FOV. Scale bar: 20-µm. (J) Ratios between FOV and illumination area as a function of light sheet lengths and widths. Blue line: PEARLS. Other lines: LLS with different widths. (D–G) show deconvolved images.

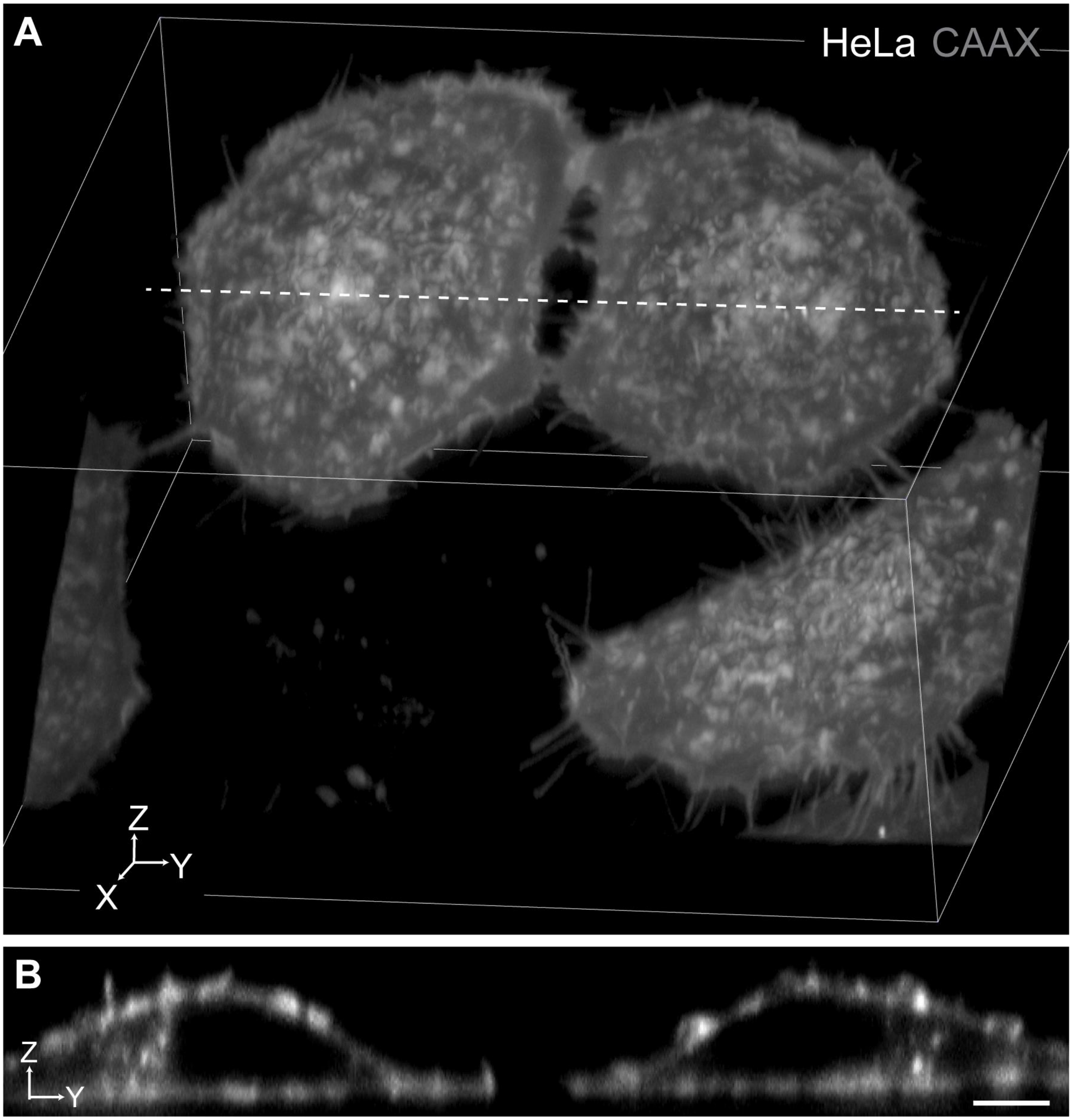


**Figure S10 | Raw data corresponding to Figure 3A,B.** (A) Volume rendering of the same field of view and time point shown in Figure 3A, displayed without deconvolution. (B) YZ orthogonal view along the same dashed line as in Figure 3A. Scale bar: 5-µm.

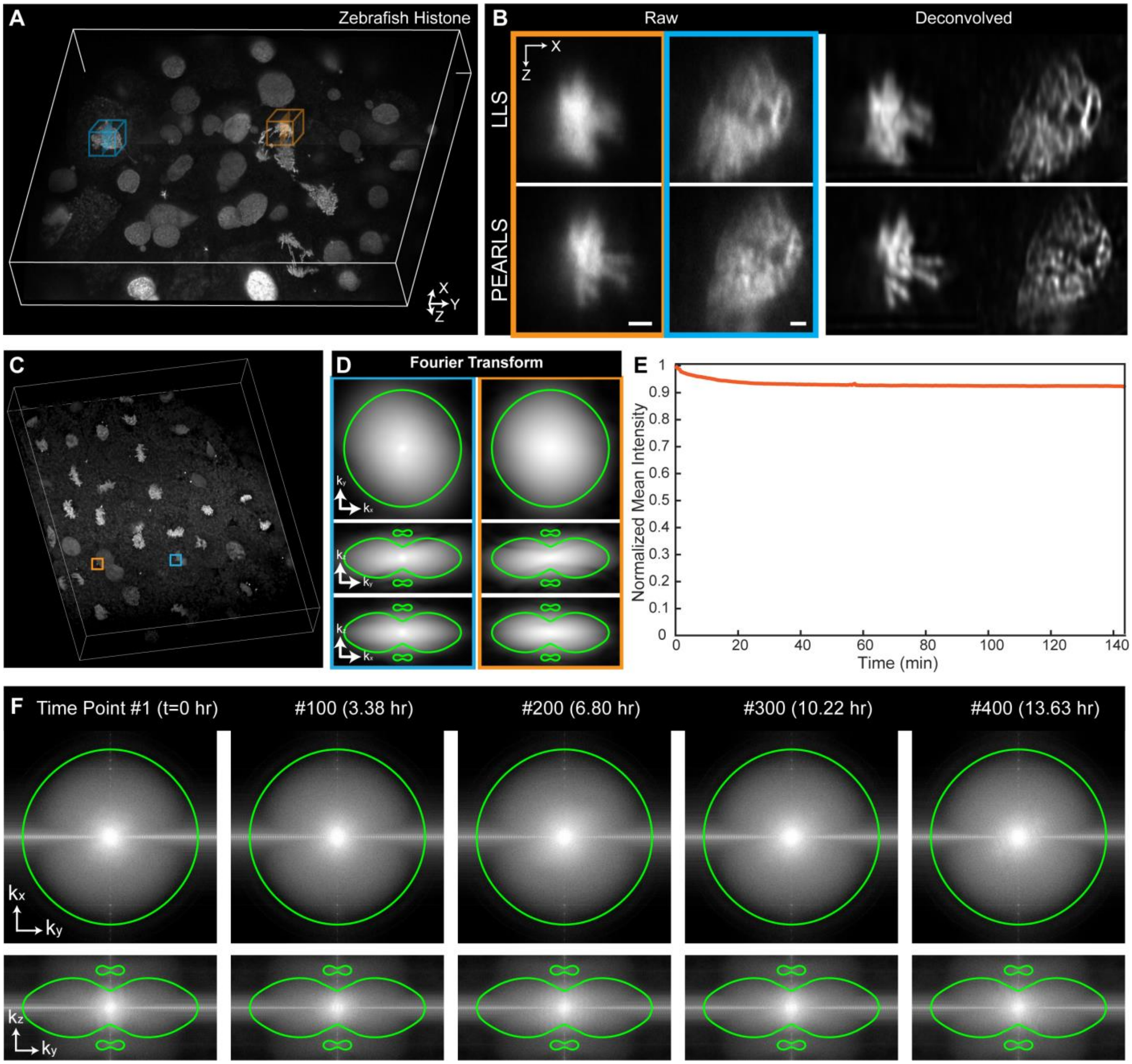


**Figure S11 | Characterization of volumetric imaging in multicellular organisms.** (A) Volume rendering of a pan-nuclear (H2B-GFP) zebrafish embryo (~4hpf) imaged using PEARLS. Bounding box: 146-µm by 178-µm by 28-µm. (B) Axial slices of the representative regions highlighted in (A), acquired using LLS (top) and PEARLS (bottom), shown as raw data (left) and after deconvolution (right). Scale bars: 2-µm. (C) Three-dimensional rendering of the volumetric dataset corresponding to Figure 4A. Representative bright puncta (colored boxes) were selected for quantitative analysis. (D) Three-dimensional Fourier transform computed from the selected regions in (C). The green contour indicates the theoretical diffraction-limit boundary defined by the simulated overall OTF at a 1% normalized amplitude threshold. (E) Mean intensity of the 3D raw data corresponding to Figure 4A over 322 time points normalized to the first time point. (F) Fourier analysis of the 3D deconvolved dataset corresponding to Figure 4C,D. Representative lateral ($k_x$-$k_y$ top row) and axial ($k_z$-$k_y$, bottom row) center slices of the three-dimensional Fourier transform at selected time points (#1, #100, #200, #300, and #400). The green contours indicate the diffraction-limited boundary defined by the simulated system OTF at a 1% normalized amplitude threshold.

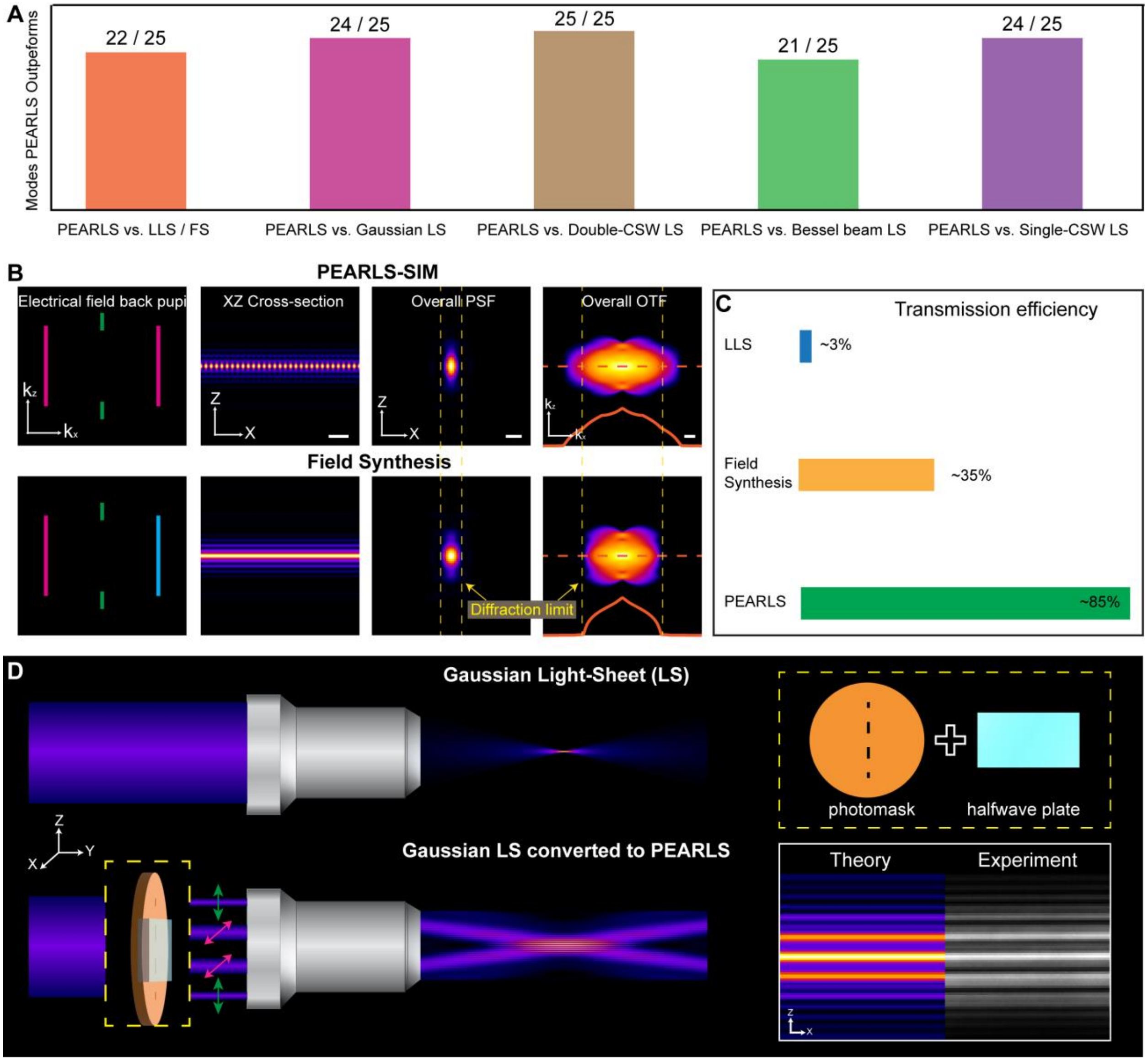


**Figure S12 | PEARLS in comparison with state-of-the-art light sheets.** (A) Strehl ratio comparisons between PEARLS and other light sheets under 25 different Zernike aberration modes. Data and color coding are consistent with Figure 1F and Figure S6. (B) Comparison of PEARLS-SIM (top) and Field Synthesis (bottom). From left to right: electrical fields at the back pupil, XZ cross-sectional illumination profiles, overall PSFs, and overall OTFs. PEARLS can generate lateral interference patterns that enable structured-illumination-based super-resolution imaging, whereas Field Synthesis is limited to diffraction-limited performance. Green, magenta, and blue denote incoherent beamlets. Yellow dashed lines indicate the diffraction limit. Orange curves show linecuts along the orange dashed line. Scale bars: 5-$\lambda_{exc}$/n (left), 1-$\lambda_{exc}$/n (middle), and 1-n/$\lambda_{exc}$ (right). $NA_{max}/NA_{min}$=0.6/0.4. (C) Transmission efficiency of the beam-shaping units for LLS (blue), Field Synthesis (orange), and PEARLS (green). (D) Conversion of a Gaussian LS (top) into PEARLS (bottom) using a single compact add-on module composed of a photomask and a HWP. Green and magenta arrows indicate two orthogonal polarization states.

### Legends for Supplementary Videos

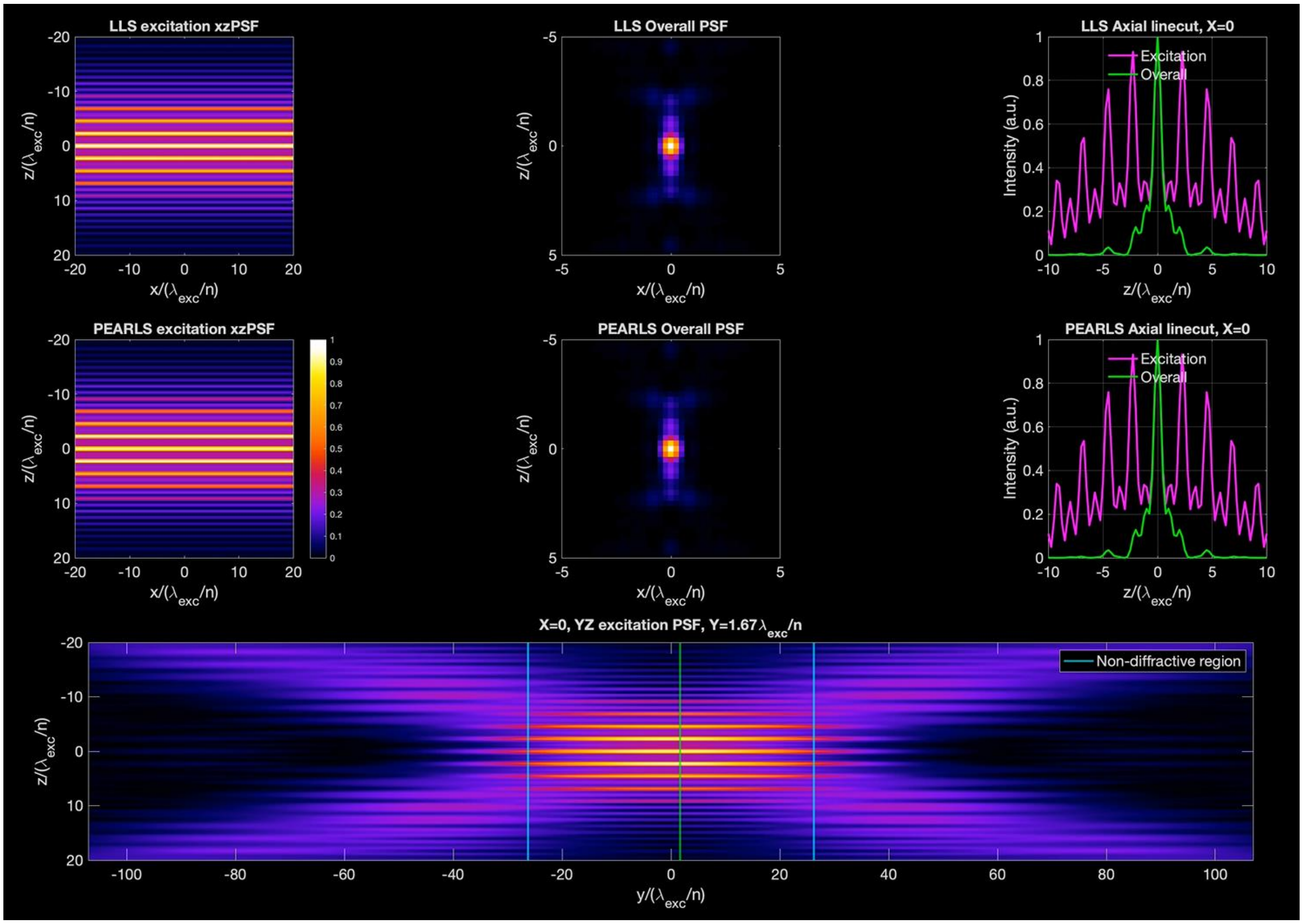


**Video S1 | Beam profiles and PSFs comparisons between dithered LLS and PEARLS.** Comparisons of LLS (top row) and PEARLS (middle row) transverse illumination profiles (left column), overall XZ PSFs (middle column), and the axial line cuts (right column) of the illumination profile (magenta) and overall PSFs (green) as a function of the propagation distance y indicated by the green line on the bottom panel that shows the longitudinal illumination profile. See also Figure 1B.

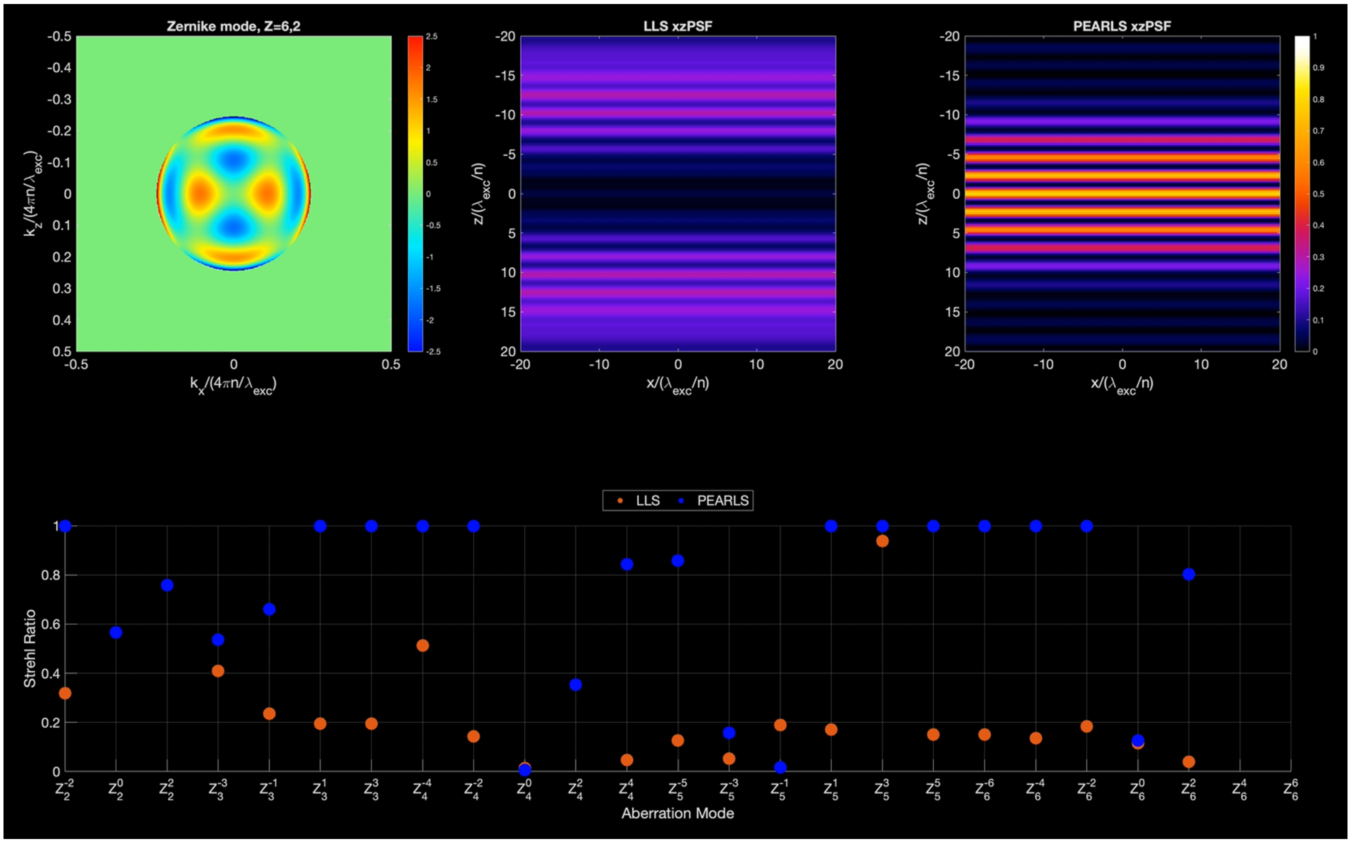


**Video S2 | Comparisons of LLS and PEARLS under aberrations.** Aberrations represented by different Zernike modes (top right panel) were applied to the rear pupil plane of LLS and PEARLS, leading to distortions of the XZ cross-section profiles of LLS (top middle panel) and PEARLS (top right panel). Bottom panel shows the Strehl ratios of LLS (orange) and PEARLS (blue) under these Zernike aberrations. See also Figure 1F.

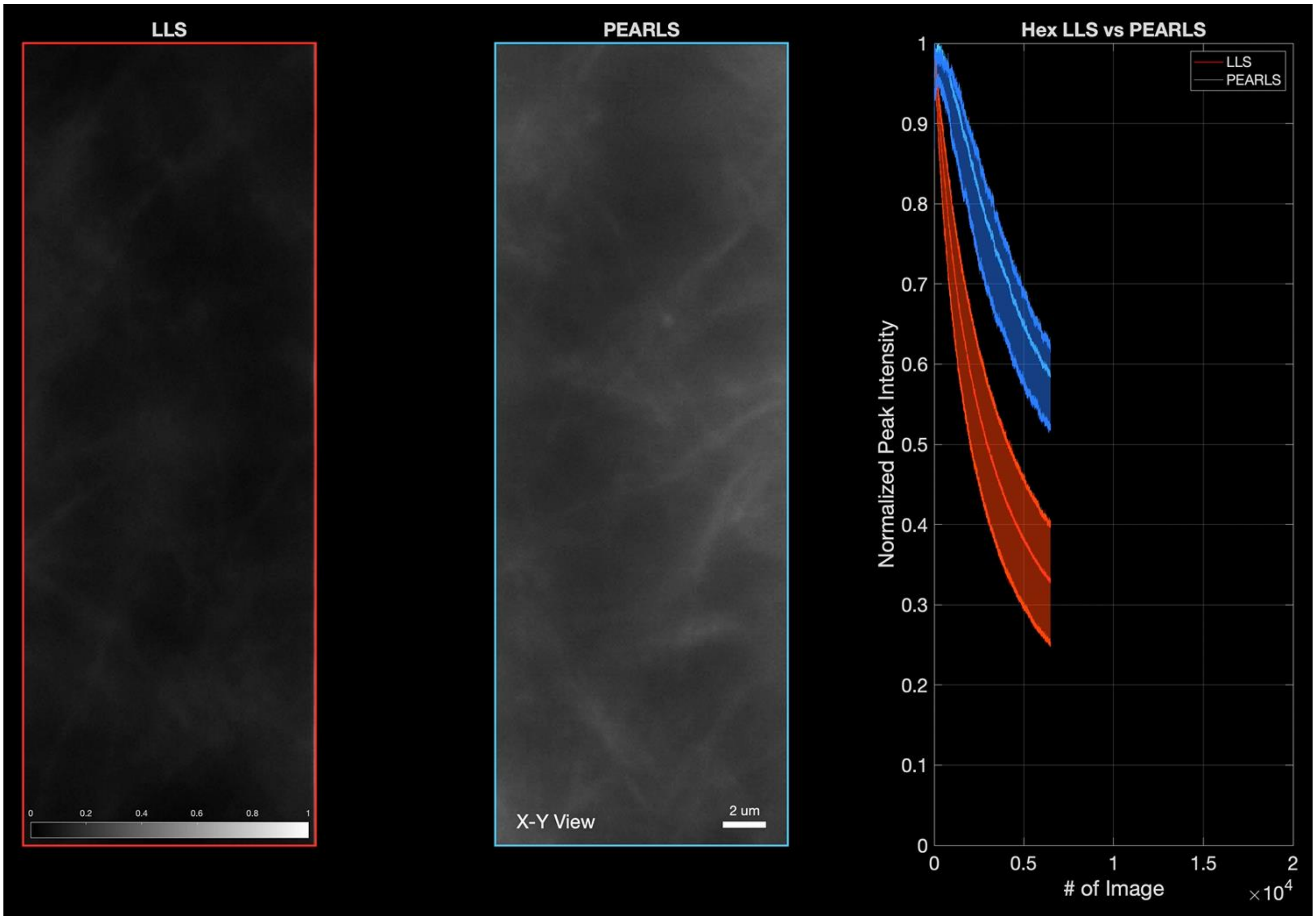


**Video S3 | Photobleaching comparisons of hexagonal type LLS and PEARLS with identical profile.** Each frame of LLS (left panel) and PEARLS (middle panel) was normalized with respect to (w.r.t.) the maximum intensity of the Frame 1 of corresponding light sheets. Right panel shows the averaged photobleaching curves of the PEARLS (blue) and LLS (red). The shaded areas indicate ±1 standard deviation (SD) from five datasets taken at five different regions of the same collagen gel. Intensities were normalized w.r.t. the maximum value of the corresponding Frame 1. See also Figure 2E&F.

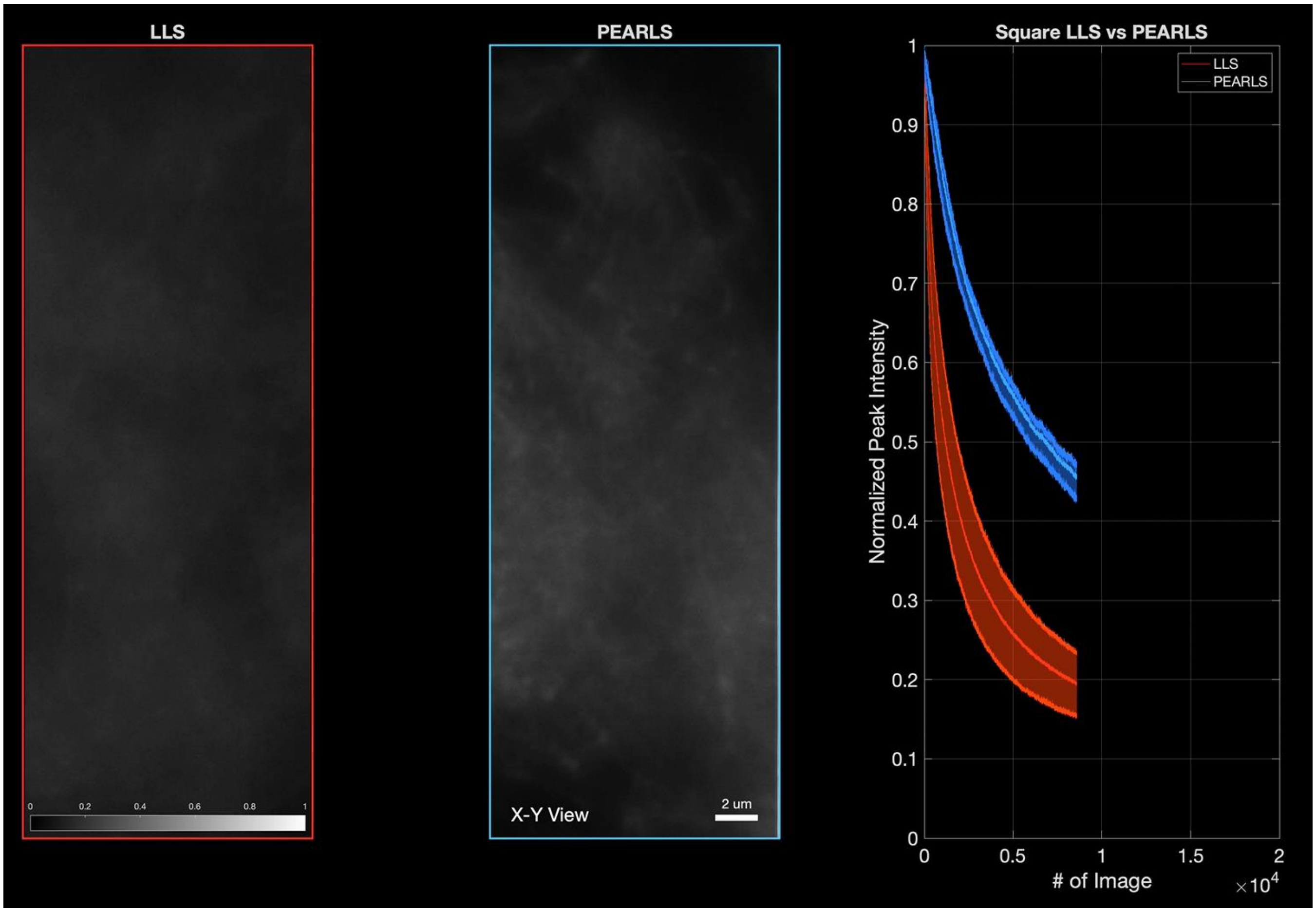


**Video S4 | Photobleaching comparisons of square type LLS and PEARLS with identical profile.** Each frame of LLS (left panel) and PEARLS (middle panel) was normalized with respect to (w.r.t.) the maximum intensity of the Frame 1 of corresponding light sheets. Right panel shows the averaged photobleaching curves of the PEARLS (blue) and LLS (red). The shaded areas indicate ±1 standard deviation (SD) from five datasets taken at five different regions of the same collagen gel. Intensities were normalized w.r.t. the maximum value of the corresponding Frame 1. See also Figure S9H.

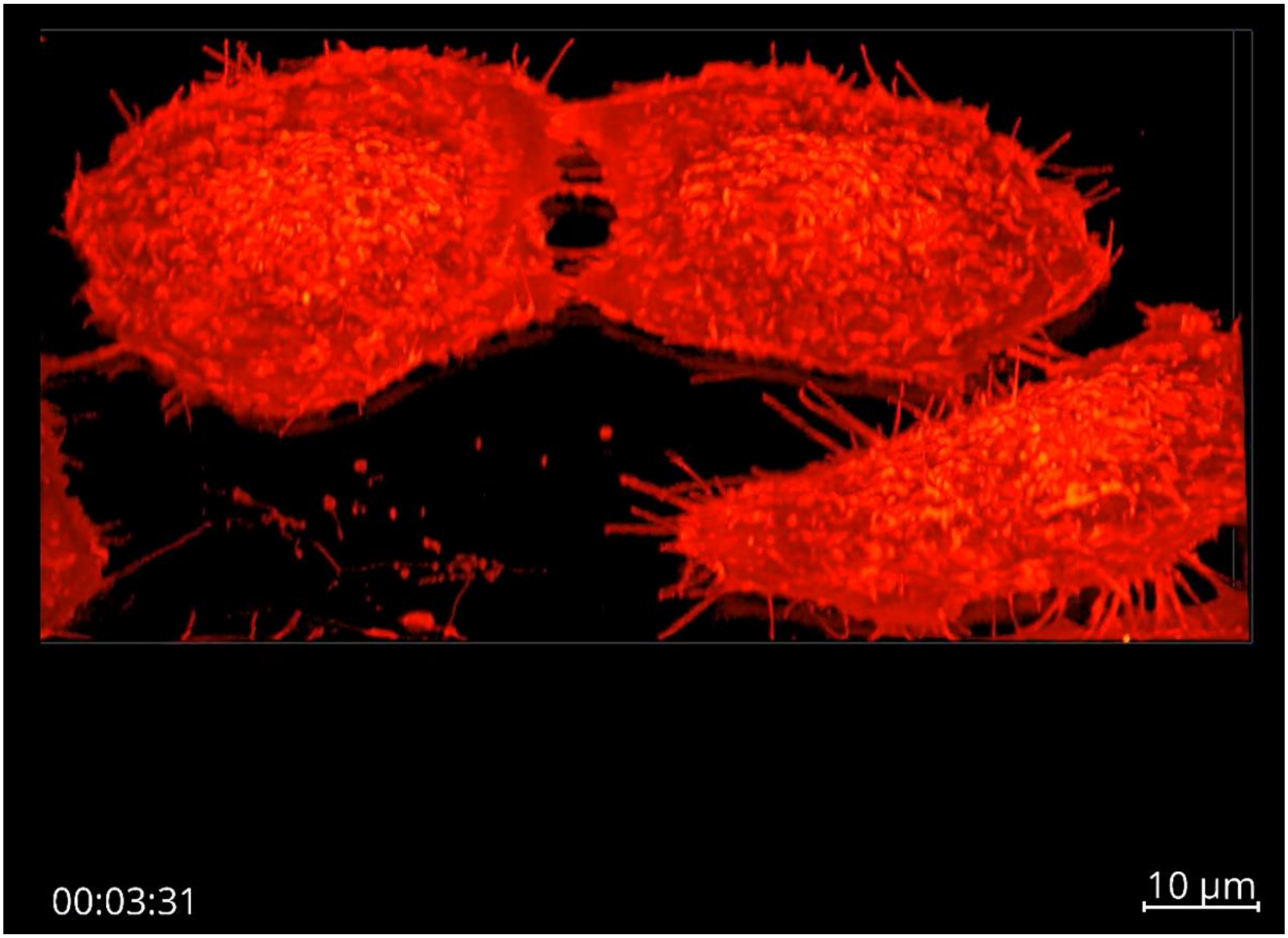


**Video S5 | Volume rendering of HeLa cells membrane dynamics.** The HeLa cells stably expresses mCherry-CAAX**.** The dataset covers 300 time points acquired at 13.2-s intervals using a sample scan mode. Bounding box: 111-μm by 110-μm by 100-μm. See also Figure 3A-C.

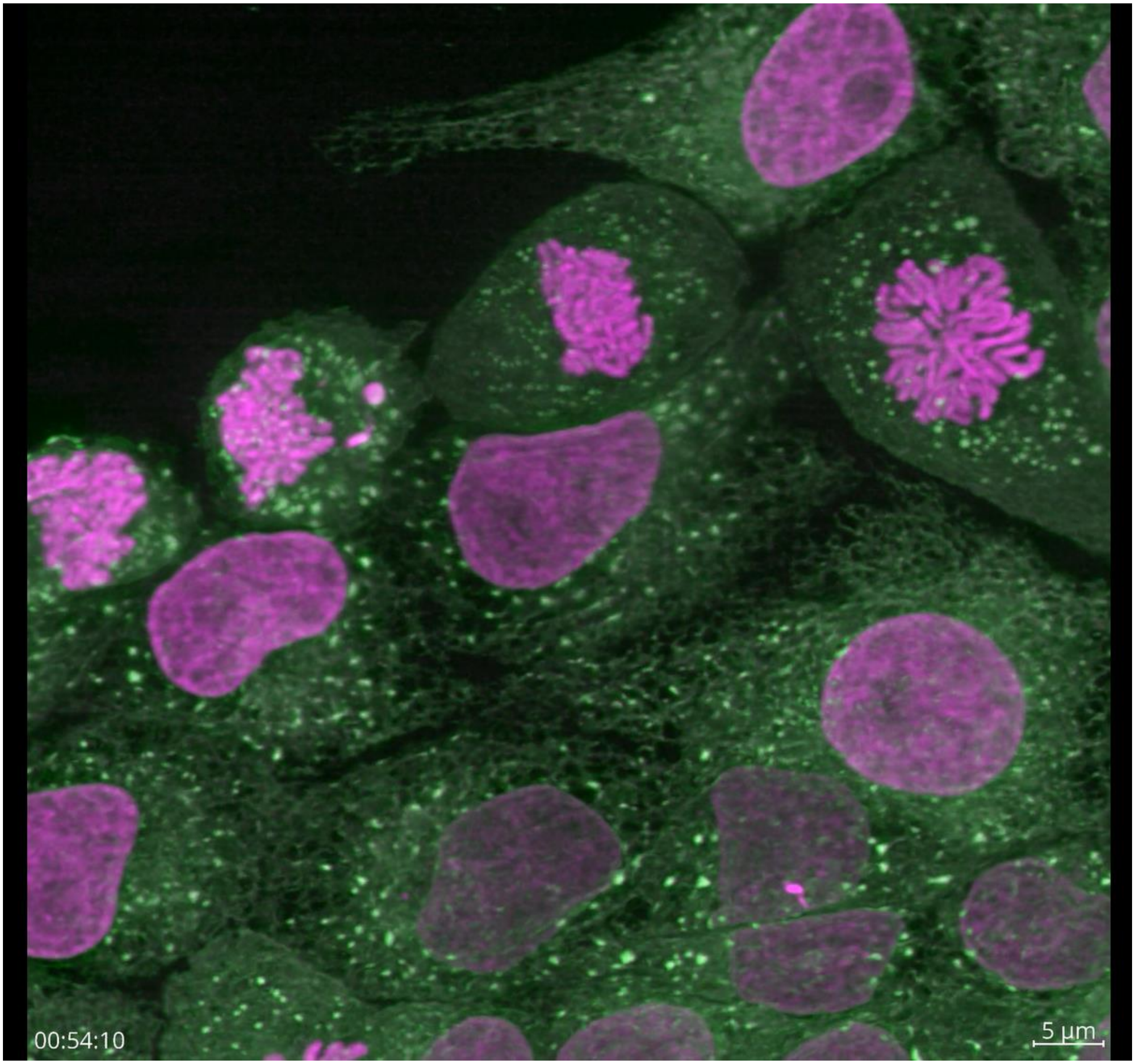


**Video S6 | Volume rendering of simultaneous two-color imaging of mitotic LLCPK1 cells.** The LLCPK1 cells stably expresses mEmerald-ER (green) and mCherry-H2B (magenta)**.** The dataset covers 400 time points acquired at 15.7-s intervals using a sample scan mode. Bounding box: 111-µm by 110-µm by 100-µm. See also Figure 3D-F.

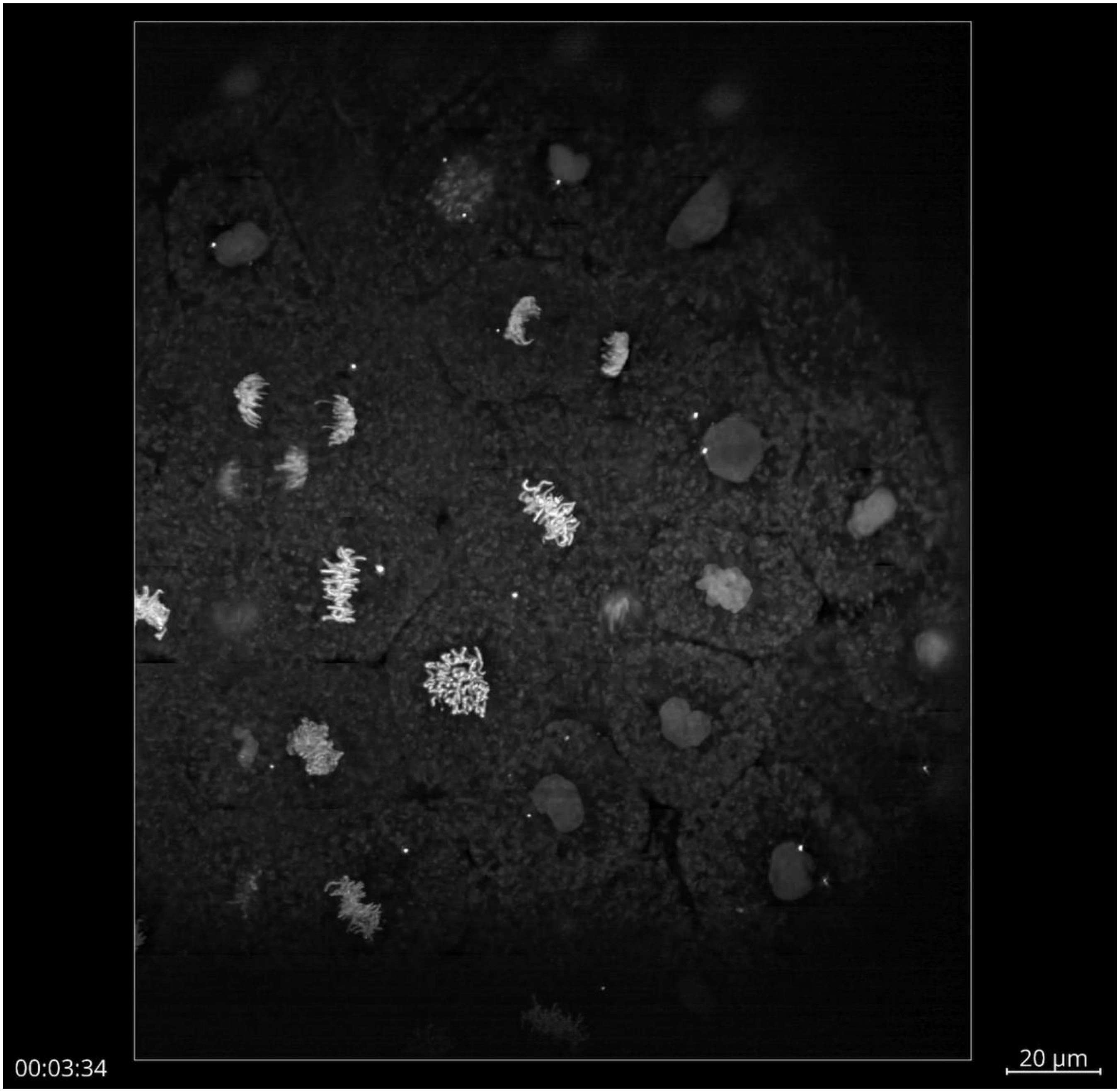


**Video S7 | Volume rendering of mitosis in zebrafish embryogenesis.** A pan-nuclear (H2A-GFP) zebrafish embryo (~3hpf) was imaged with a total of 322 time points at 26.8-s time intervals. Bounding box: 264-µm by 221-µm by 60-µm. See also Figure 4A,B.

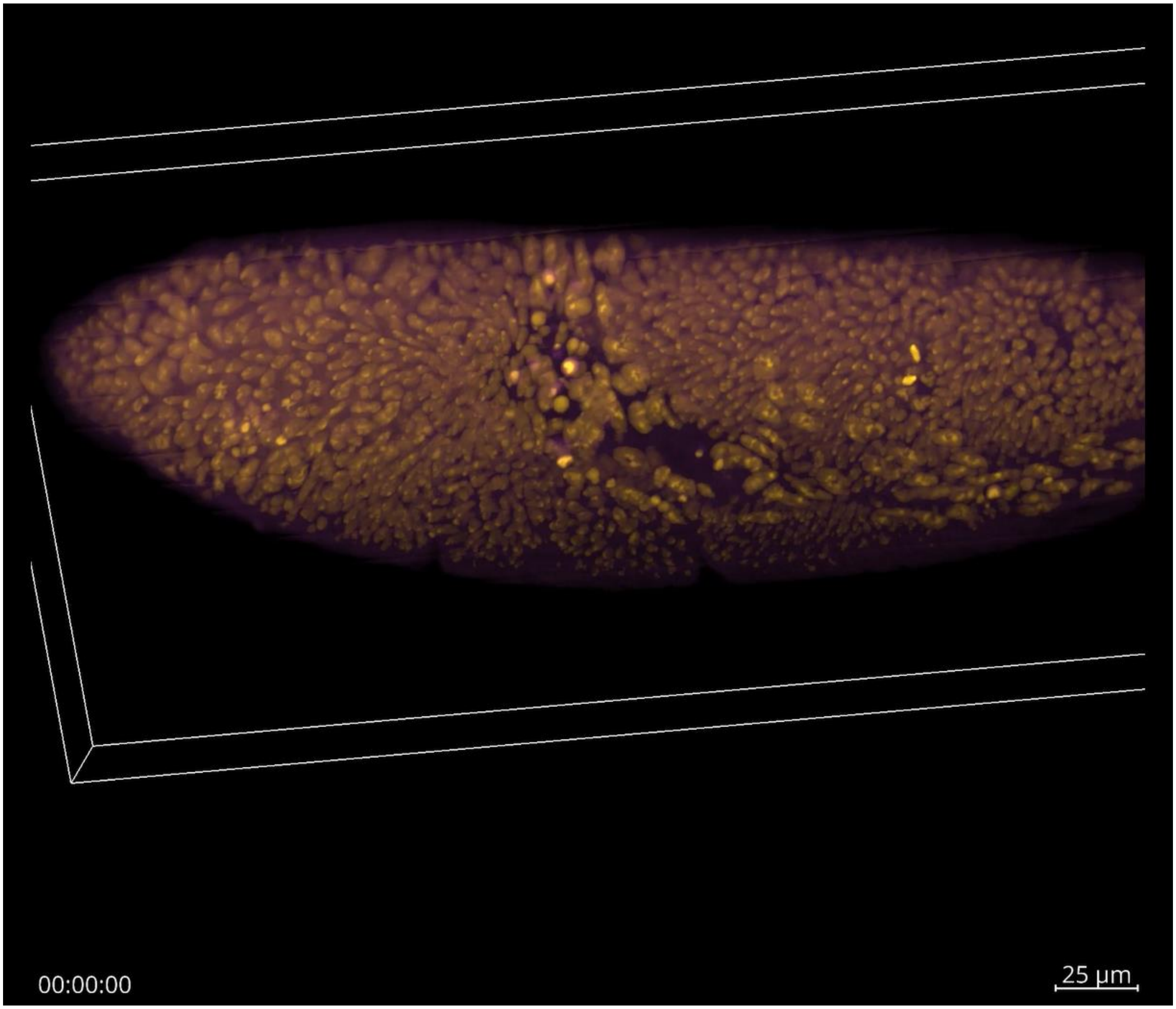


**Video S8 | Volume rendering of large-scale cell migration in Drosophila development.** A *Drosophila melanogaster* embryo expressing mNeonGreen-MCP (magenta) and mRFP-histone (yellow) was imaged with a total of 412 time points at 2-min time intervals. Bounding box: 432-µm by 194-µm by 60-µm. See also Figure 4C,D.
